# Single-Cell RNA and ATAC Sequencing Reveal Cellular Heterogeneity and Chromatin Accessibility Dynamics in Young Chinese Breast Cancer Patients across Pre- and Post-Neoadjuvant Therapy

**DOI:** 10.1101/2025.10.14.682233

**Authors:** Xue Han, Bin Wang, Wenting Xiang, Fan Lin, Bo Sun, Beibei Yang, Zhihan Ruan, Shasha Liu, Shuo Li, Hong Liu, Jian Liu

## Abstract

Young breast cancer (YBC) patients (age ≤ 40) exhibit aggressive features and poor prognosis, while facing complex needs including breast-conserving surgery and fertility preservation. Neoadjuvant therapy (NAT) is important for tumor downstaging and breast-conserving surgery in YBC treatment, but the NAT response mechanisms in YBC patients remain unelucidated. Here, we analyzed pre- and post-NAT samples from Chinese YBC patients using scRNA-seq and scATAC-seq. We found that NAT reshaped cellular heterogeneity in the tumor microenvironment (TME), reducing epithelial and T cells while increasing endothelial cells and fibroblasts. Residual cancer cells showed enriched epithelial-mesenchymal transition (EMT) and stromal remodeling programs. NAT responders had fewer luminal-like cells and retained basal-like cells in an early hybrid EMT state, whereas non-responders maintained both populations with late hybrid EMT. We identified therapy-resistant genes and motifs (e.g., VTCN1, PROM1, MZB1, Fox family) and linked CXCL13 upregulation to poor NAT response and tumor-specific T-cell expansion. Cell–cell communication analysis revealed NAT reprograms the TME by suppressing VEGF and TNF signaling in basal cells, while residual luminal cells transmit EGFR and CD47–SIRPA signals.

## INTRODUCTION

Breast cancer is the most commonly diagnosed cancer among women worldwide and the leading cause of cancer death[1–2]. The proportion of young women (typically defined as those diagnosed at age ≤ 40) among breast cancer patients has shown an increasing trend in recent years[3–4]. The average age at diagnosis for breast cancer in China is younger than that in many Western countries [5–6]. Young patients with breast cancer usually exhibit aggressive features, poor histological grade, and unfavourable prognosis[7–8]. Beyond undergoing anti-cancer therapy, they have to navigate complex trade-offs concerning their desire for breast conservation and the need for fertility preservation [9]. Neoadjuvant therapy is an important strategy to reduce tumor size, potentially turning initially inoperable tumors into ones that can be removed surgically, thus increasing breast conservation rates, and it also allows for dynamic assessment of how the tumor responds to treatment [10–11]. Investigating the response patterns to NAT and their heterogeneity within YBC patients is important for optimizing treatment strategies and enhancing quality of life.

Currently, some research has explored NAT for breast cancer through multi-omics analyses. A study has provided multi-omics molecular profiles of responders and non-responders in Chinese breast cancer patients, covering ER−HER2+, ER+HER2+, and ER−HER2− subtypes[12]. This study was based on bulk sequencing technology, which made it difficult to effectively characterize the specific changes in TME and cancer cell heterogeneity before and after neoadjuvant therapy. Several recent studies used scRNA-seq or scATAC-seq to elucidate breast cancer microenvironments [13–16]. A scRNA-seq study of triple-negative breast cancer during neoadjuvant chemotherapy (NAC) demonstrated that transcriptional profiles remained stable in the NAC-response group. In contrast, the non-response group exhibited dysregulation in genes associated with immune cell proliferation and differentiation[13]. Another study identified nine distinct epigenetically-regulated cancer cell states in tamoxifen-resistant breast cancer[14]. However, neoadjuvant treatment (NAT) response targeting YBC patients in single-cell resolution remains unelucidated, especially for the underlying molecular mechanisms governing treatment response heterogeneity between good- and poor-NAT response patients. Integrating scRNA-seq and scATAC-seq to elucidate cellular heterogeneity and chromatin accessibility dynamics in YBC patients across pre- and post-NAT could offer promising insights into treatment-induced tumor microenvironment remodeling and therapeutic resistance mechanisms.

Here, we profiled 29 tumor and non-tumor samples from 15 young Chinese women with breast cancer, pre-, post-, and non-NAT, employing scRNA-seq and scATAC-seq to characterize the treatment-induced dynamic remodeling of the TME and cancer cell heterogeneity. The analysis revealed significant changes in TME cellular composition following neoadjuvant therapy, characterized by reduced abundance of epithelial cells, T cells, and myeloid cells, coupled with increased abundance of endothelial cells, fibroblasts, and perivascular-like (PVL) cells. We revealed that residual cancer cells after NAT were enriched for genes and motifs associated with epithelial-mesenchymal transition (EMT) and stromal remodeling, suggesting this may constitute an important mechanism underlying treatment resistance. We observed that samples with better pathological response (grade Ib–IIIb) exhibited preferential reduction of luminal-like cancer cells with retention of basal-like cells, whereas those with poorer response (grade Ia) maintained both subtypes. After NAT, tumor cells in good-response samples mainly exhibited an early hybrid EMT state, whereas in poor-response samples, the residual tumor cells were mainly in a late hybrid EMT state. Consistently, we identified chromatin accessibility dynamics of therapy-resistant motifs, potentially unveiling core mechanisms of treatment resistance in YBC. Moreover, elevated CXCL13 expression in terminally exhausted T cells (tTex) suggests its potential as a biomarker for poor prognosis. In post-NAT samples, basal cells tend to adopt a matrix-dependent and differentiation/repair-oriented adaptation, whereas luminal cells shift toward proliferation and immune evasion; key signaling axes, including Integrin, VEGF, and EGFR, collectively shape therapeutic response differences and may constitute actionable targets for combinatorial intervention. In brief, this work enhances our understanding of the molecular mechanisms underlying neoadjuvant therapy response in young breast cancer patients.

## RESULTS

### Overview of Integrated Single-Cell Multi-omics in Breast Cancer Pre- and Post-Neoadjuvant Therapy

To investigate cellular heterogeneity and epigenetic regulation in young breast cancer patients before and after neoadjuvant therapy, we enrolled 15 female patients aged 40 years or younger. Among them, ten patients met the clinical indications for neoadjuvant therapy. They underwent surgical resection following treatment, with tumor tissues collected either at the time of core needle biopsy or surgery. The remaining five patients, who did not meet neoadjuvant therapy criteria, received upfront surgery and served as untreated controls. Single-cell RNA sequencing (scRNA-seq) and/or single-cell assay for transposase-accessible chromatin using sequencing (scATAC-seq) were performed to profile tumor and control tissues at single-cell resolution. Based on immunohistochemistry (IHC) subtyping, the cohort included six HR+HER2-, four HR+HER2+, one HR-HER2+, and four triple-negative breast cancer (TNBC) cases (Tables S1 and S2). In total, 29 tissue samples were collected, comprising 24 tumor samples and five non-tumor control (NT) samples. Among these, six tumor samples were obtained pre-NAT via biopsy, and 23 were collected post-NAT during surgery. For library preparation, two samples were processed using the 10x Genomics 3’ scRNA-seq platform, while 16 were profiled using the 5’ scRNA-seq platform. Additionally, 11 samples were subjected to scATAC-seq analysis (Table S2). All 5’ scRNA-seq libraries were further enriched with immune repertoire profiling to capture both B cell receptor (BCR) and T cell receptor (TCR) sequences, enabling a comprehensive analysis of the tumor immune landscape (Fig. 1A).

**Figure 1.**
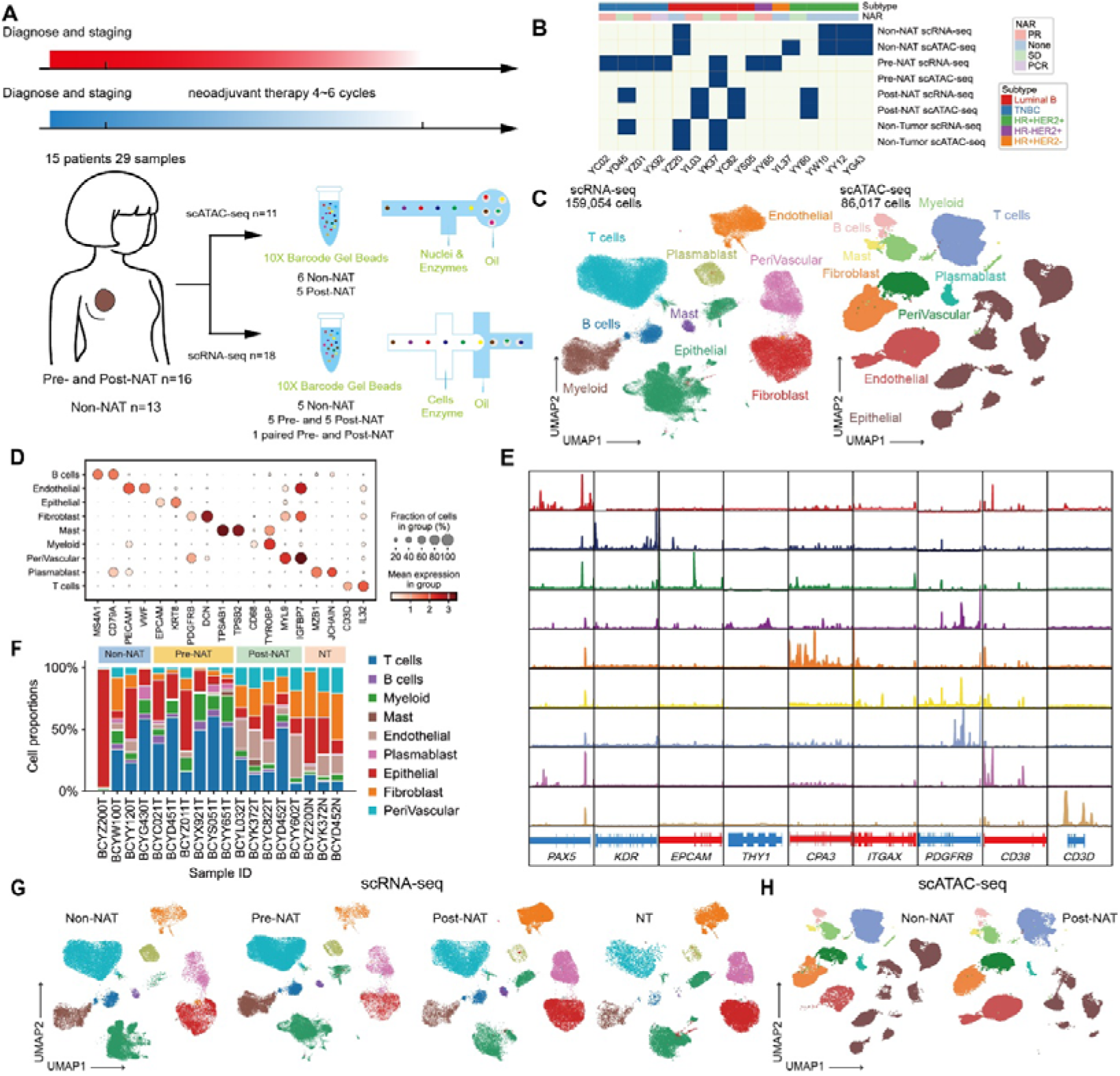
Overview of Single-Cell Multi-omics data in YBC Patients across Pre- and Post-NAT. (A) An overview of the study design. (B) Heatmap shows the details of sample origin, treatment regimen, intrinsic subtype, and NAT response (NAR) evaluation. (C) UMAP visualization of scRNA-seq (left) and scATAC-seq (right) cell clusters. Each dot represents a cell, colored by its cell type. (D) The dotplot heatmap shows the marker genes of each cell type in scRNA-seq data. (E) ATAC-seq aggregated signal tracks illustrate peaks of chromatin accessibility around the approximate transcription start sites of each cluster’s marker genes. Red coloring denotes positive-strand genes, with blue coloring for negative-strand genes. (F) Proportions of cell types across tumors in scRNA-seq data. (G) UMAP visualization of scRNA-seq data of cell types in each condition, including non-NAT, pre-NAT, post-NAT, and non tumoral samples (NT). (H) UMAP visualization of scRNA-seq data in each condition, including pre-NAT and post-NAT.

To evaluate the impact of neoadjuvant therapy on breast cancer, we constructed a comprehensive single-cell RNA sequencing (scRNA-seq) atlas encompassing tumor samples across various treatment states. This atlas included four untreated samples, five samples collected before NAT, four post-NAT samples, one matched pre- and post-NAT pair, and three non-tumor control (NT) samples (Fig. 1B). Following stringent quality control (QC) filtering, we retained a total of 159,054 high-quality cells for downstream analysis (Fig. 1C and Supplementary Fig. 1A and 1C). To further investigate the epigenetic alterations associated with neoadjuvant therapy, we generated a single-cell chromatin accessibility (scATAC-seq) atlas, comprising five untreated tumor samples, four post-NAT samples, and two NT samples (Fig. 1B). A total of 86,017 high-quality cells were obtained from these samples and used in subsequent analyses (Fig. 1C and Supplementary Fig. 1B and 1D). Based on established marker genes, we identified nine major cell types across both modalities: epithelial cells, endothelial cells, B cells, T cells, fibroblasts, myeloid cells, plasma cells, perivascular-like (PVL) cells, and mast cells (Fig. 1D-E and Supplementary Fig. 1E-F). All nine cell types were consistently detected across the 29 samples, indicating their broad presence within breast tissue. Compared to NT samples, tumor samples displayed markedly increased cellular heterogeneity. While NT samples showed a relatively stable cellular composition dominated by epithelial, endothelial, fibroblast, and PVL cells, tumor samples exhibited greater complexity, with notable enrichment of infiltrating T cells, B cells, and antigen-presenting cells (Fig. 1F). Interestingly, in the uncorrected UMAP[17] embeddings, immune-related cell types displayed relatively consistent distributions across samples, whereas epithelial cells exhibited pronounced inter-patient heterogeneity. This pattern was observed in both scRNA-seq and scATAC-seq datasets, highlighting the multilayered molecular diversity of epithelial cells across tumors (Fig. 1G and 1H).

Cellular composition analysis revealed a marked reduction in the proportions of epithelial and immune cells in post-NAT samples compared to both pre-NAT and untreated controls. In contrast, endothelial and stromal cell populations exhibited a notable expansion. To validate these observations, we employed the miloR[18] to perform differential abundance analysis across cell subpopulations. Consistent with our initial findings, the analysis demonstrated a significant decrease in epithelial cells, T cells, and myeloid cells, accompanied by a pronounced enrichment of endothelial cells, fibroblasts, and PVL cells in post-NAT tumors(Supplementary Fig. 1G). These shifts in cellular abundance suggest that, while neoadjuvant therapy (NAT) effectively eliminates tumor cells, it concurrently depletes tumor-infiltrating immune populations, potentially impairing anti-tumor immune responses. Concurrently, the expansion of endothelial and stromal compartments highlights a profound remodeling of the tumor microenvironment (TME) following treatment. This reorganization likely encompasses angiogenic reprogramming and stromal activation, which may collectively foster a more immunosuppressive milieu and influence post-NAT immune surveillance and relapse risk. Previous studies have reported that NAT can attenuate tumor-associated immune infiltration while promoting the proliferation of stromal elements, including fibroblasts and endothelial cells[19]. These dynamics may be driven by therapy-induced inflammatory responses, tissue repair mechanisms, and survival adaptations of residual tumor cells. As such, elucidating the mechanisms underlying therapy-induced TME remodeling is critical for refining subsequent immunotherapeutic strategies and rationalizing combinatorial treatment approaches.

### Characterization of Tumor Cells and Their Dynamical Changes Across NAT

We re-clustered 38,680 epithelial cells in scRNA-seq data and 29,719 epithelial cells in scATAC-seq data, and then annotated major lineages: a basal lineage (marked by KRT5, KRT14, and ACTA2), a hormone receptor–associated luminal lineage (LumHR, marked by KRT8, KRT18, and KRT19), and a secretory luminal lineage (LumSec, expressing KRT15 and SCGB2A2) (Fig. 2A and 2B). These lineages were each initially classified into normal type and cancer type based on sample origin. To further distinguish between tumor and normal epithelial cells, we applied inferCNV[20] and a scATAC-seq–based copy number variation (CNV) inference method[21], with immune cells used as references. In the scRNA-seq data, epithelial cells from non-tumor samples and tumor-derived cells with low CNV scores were classified as normal epithelial cells, whereas those with high CNV scores were designated as malignant epithelial cells. We generated a UMAP projection of the CNV score heatmap and found a clear separation of tumor cells, while normal epithelial cells from different samples clustered together. (Fig 2C). Projection of transcriptome-based annotations onto scATAC-seq data yielded highly consistent cell-type identities and proportions across samples (Fig. 2A and 2B). According to the annotated epithelial cells, we observed heterogeneity in epithelial cell composition among different patient groups in both scRNA-seq and scATAC-seq data (Fig. 2B).

**Figure 2.**
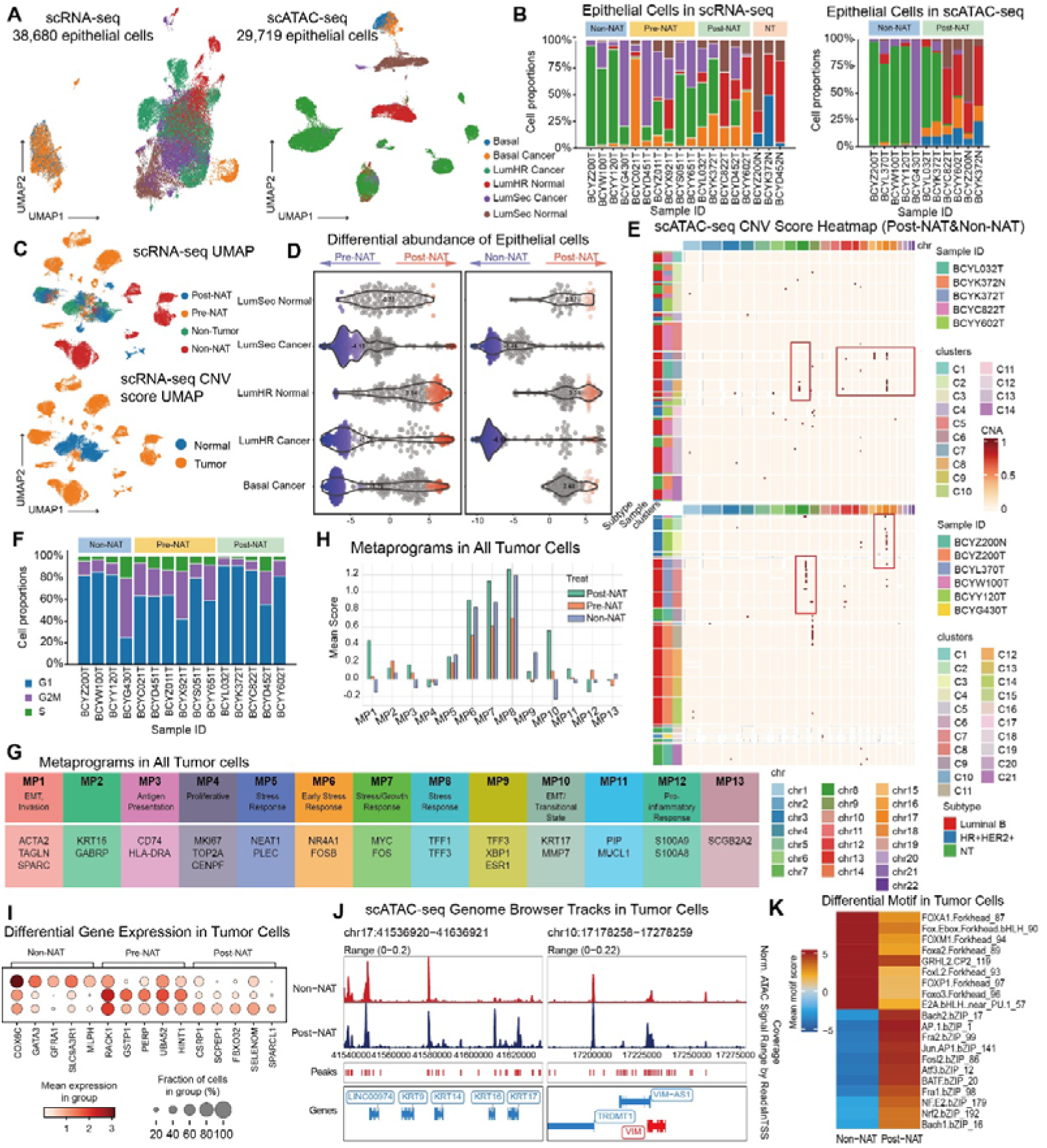
Characterization of tumor cells and their dynamical changes across NAT. (A) UMAP visualization of epithelial cells in scRNA-seq data (left) and scATAC-seq data (right). (B) Proportions of epithelial subtypes across samples in scRNA-seq data (left) and scATAC-seq data (right). (C) UMAP visualization of epithelial cells from tumor samples in scRNA-seq data according to the CNV alteration heatmap, colored by intrinsic subtypes (top) and CNV status (bottom). (D) Beeswarm plot of the log fold change distribution across epithelial cells from pre-, post- and non-NAT samples in scRNA-seq data. DA neighborhoods at FDR 10% are colored. (E) CNV score heatmap of epithelial cells in scATAC-seq data. (F) Proportions of cell-cycle states across samples in scRNA-seq data. (G) Meta programs (MPs) detected from all tumor cells using scRNA-seq data. (H) Barplot shows the enrichment score of MPs in non-NAT, pre-NAT, and post-NAT groups. (I) Dotplot heatmap shows the differential expression genes of tumor cells in non-NAT, pre-NAT, and post-NAT. (J) Pseudobulk scATAC-seq signal tracks illustrating differences in chromatin accessibility profiles around the transcription start sites (TSS) of representative EMT marker genes between post-NAT and non-NAT samples. (K) Heatmap shows differential motifs of tumor cells between non- and post-NAT samples.

Next, we investigated the changes in epithelial cell compositions pre-, post-, and non-NAT. Both transcriptomic and epigenomic analyses revealed lower mean CNV scores in post-NAT samples, supporting the notion that NAT mitigates genomic instability in tumor cells (Supplementary Fig. 2A). Abundance analysis revealed a significant reduction of luminal malignant epithelial cells in post-NAT samples compared to pre-NAT and non-NAT samples (Fig. 2D). Basal cancer cells were relatively enriched in pre-NAT biopsies, markedly reduced after therapy, and modestly increased compared to non-NAT samples. For normal epithelial cells, LumHR cells were significantly enriched in post-NAT tumors, whereas LumSec cells showed fluctuating abundance. Notably, the post-NAT cohort predominantly consisted of HR+/HER2-subtype (Table S1). Consistent with previous research, NAT exerted stronger cytotoxic effects on highly proliferative breast cancer subtypes, such as TNBC and HER2+ breast cancer; the HR+/HER2− subtype, which was largely composed of low-proliferative LumHR cancer cells, tends to show less responsiveness to NAT[22]. Thus, our analyses indicated that in young patients with breast cancer, NAT may preferentially eliminate chemosensitive tumor cells, resulting in the relative enrichment of residual LumHR cells following treatment.

Next, we examined chromosomal CNV alterations in tumor cells across tumor groups. CNV inference from scRNA-seq revealed that, compared with adjacent normal tissue, pre-NAT samples displayed focal CNV alterations, including deletions on chromosomes 3, 5, 13, and 14, and amplification of chromosome 18. (Fig. 2C and Supplementary Fig. 2B). Tumor cells from non-NAT samples exhibited widespread CNV abnormalities, with deletions on chromosomes 5 and 22, and extensive amplifications across chromosomes 1, 2, 7–10, 15–20, and X. In contrast, tumor cells from post-NAT samples showed substantially attenuated CNV profiles, with amplifications restricted to chromosomes 8 and 20 and no significant deletions (Supplementary Fig. 2B). The CNV patterns inferred from scATAC-seq were concordant with those in scRNA-seq data. Specifically, tumor cells from non-NAT samples exhibited prominent CNV changes on chromosomes 1, 8, and 17, as well as additional chromosomal alterations; post-NAT tumor cells showed limited abnormalities, which were mainly concentrated on chromosomes 8, 12, and 15 (Fig. 2E). Collectively, these findings indicate that in YBC patients, NAT exerts a genome-stabilizing effect: while non-NAT samples display the most genomic alterations, post-NAT samples exhibit CNV landscapes trending toward normal.

To further clarify how NAT impacted biological processes in tumor cells of YBC patients, we performed cell cycle state and pathway enrichment analyses on tumor cells. Cell cycle profiling revealed a significant reduction of tumor cells in G2/M phase (reflecting active proliferation) after NAT, indicating that NAT suppressed the G1/S-to-mitosis transition and thereby curtails tumor cell proliferation (Fig. 2F). Complementary pathway enrichment analyses using gseapy[23] demonstrated the downregulation of key proliferative programs, including Cell Cycle, DNA Replication, E2F Targets, and G2-M Checkpoint [24–26] (Supplementary Fig. 2C). Pathways associated with extracellular matrix remodeling, inflammatory signaling, and survival responses were upregulated in tumor cells from post-NAT samples, such as Epithelial Mesenchymal Transition, TNF-α Signaling via NF-κB, and KRAS Signaling Up (Fig. 2F). Together, these findings suggested that NAT effectively eliminated highly proliferative tumor cells in YBC patients. Meanwhile, residual cells may activate EMT-, NF-κB-, and KRAS-associated programs to engage survival and escape mechanisms, which could be potential drivers of therapy resistance.[27–30].

We also applied GeneNMF [31] to infer meta-programs (MPs) and the distribution of MP scores across non-NAT, pre-NAT, and post-NAT samples (Fig. 2G and Supplementary Fig. 2D). We identified 13 functional MPs and found their significant variation across treatment groups (Fig. 2H). In the post-NAT samples, MP1, MP10, and MP11 were relatively elevated compared with pre-NAT and non-NAT samples. These modules were enriched for genes associated with cytoskeletal organization, myofibroblast-like transition, epithelial–mesenchymal transition (EMT), and extracellular matrix (ECM) remodeling, including ACTA2, MYLK, TAGLN, and MMP7[32–36]. Such transcriptional reprogramming may represent both adaptive responses to therapy and potential resistance mechanisms. MP3, which was enriched for antigen presentation genes (CD74 and HLA-DRA), had higher scores in both pre- and post-NAT samples compared with non-NAT samples [37]. Since non-NAT samples were mainly HR+/HER2+ subtype, which were typically characterized by weak baseline immune activity[38], the low MP3 activity likely reflects their “immune-cold” phenotype rather than therapy-induced changes. MP6, MP7, MP8, MP9, and MP13 were enriched for stress response, hormone signaling, and differentiation genes, including FOS and JUN (classic immediate-early response genes), TFF1 (ER downstream target), SCGB2A2 (secretory luminal marker), and ESR1 (ERα)[39–40]. MP12 was relatively elevated in the pre-NAT group (mainly contained TNBC samples), and was enriched for genes associated with inflammatory microenvironments and myeloid infiltration (S100A8 and S100A9) [41]. This suggested that TNBC samples had stronger inflammatory activation, which may confer higher immunogenic plasticity and treatment responsiveness.

To validate the MP analysis, we further evaluated highly variable genes and motifs across the three groups. COX6C, GATA3, GFRA1, SLC9A3R1, and MLPH were highly expressed in non-NAT samples (Fig. 2I). These genes were associated with epithelial differentiation, hormone receptor signaling, and luminal-like breast cancer characteristics[42–44]. Tumor cells in non-NAT samples also showed higher accessibility of Forkhead family motifs (FOXM1, FOXO3, FOXP1, FOXL2, FOXA1), as well as GRHL2 and E2A.bHLH in scATAC-seq data, which were associated with epithelial maintenance, cell-cycle regulation, and differentiation[45–48] (Fig. 2K). Tumor cells in pre-NAT samples highly expressed genes linked to stress response (RACK1), glutathione metabolism and detoxification (GSTP1), p53/p63-mediated apoptosis and cell–cell adhesion (PERP), ribosomal fusion protein essential for translation and genome stability (UBA52), and tumor suppressor involved in transcriptional regulation and apoptosis (HINT1), reflecting a metabolically active and stress-loaded state[49–51] (Fig. 2I). Tumor cells in post-NAT samples showed high expression of CSRP1, SCPEP1, FBXO32, SELENOM, and SPARCL1, genes associated with smooth muscle–like differentiation, cytoskeletal remodeling, and EMT[52–54] (Fig. 2I). Chromatin accessibility profiling revealed greater accessibility at EMT/basal-associated loci, including VIM, KRT14, and KRT16 in post-NAT samples (Fig. 2J). Motifs of the AP-1 complex (Jun, Fra1, Fra2, Fosl2, ATF3, BATF) were enriched in post-NAT samples, indicating activation of a canonical EMT regulatory axis[55–57] (Fig. 2K). Motifs for Bach1/Bach2 and Nrf2/NF-E2 were also upregulated in post-NAT samples, reflecting activation of oxidative stress and metabolic reprogramming pathways after NAT[58–59] (Fig. 2K). These findings validated the patterns of each group: non-NAT tumors were well-differentiated and less immunogenic; pre-NAT tumors showed low activity in hormone-related pathways but high activity in stress and inflammatory modules; post-NAT tumors reflected substantial structural and functional remodeling modules.

### Heterogeneity Analysis of Neoadjuvant Therapy Response in Tumor Cells

To further dissect the heterogeneity of NAT responses among YBC patients, we performed an integrative analysis of residual tumor cells from post-NAT samples and assessed differential transcriptional programs. Based on pathological evaluation after neoadjuvant therapy (NAT), tumors were stratified into good- and poor-response groups. Specifically, BCYL032T, BCYK372T, and BCYD452T were classified as the poor-response group, with postoperative pathological grades of Ia. BCYC822T and BCYY602T, graded as Ib-IIa and IIa respectively, were classified as the good-response group (Table S1). We further applied GeneNMF to identify key transcriptional modules in post-NAT samples, which revealed 12 distinct MPs (Fig. 3A and Supplementary Fig. 2E). MP1, MP6, and MP10 were significantly enriched in the good-response group and were associated with cytoskeletal remodeling, EMT, and immune responses, indicating enhanced transcriptional plasticity and therapy sensitivity. In contrast, MP5, MP7, MP8, MP9, MP11, and MP12 were preferentially enriched in the poor-response group, involving stress response, secretory programs, heightened cell-cycle activity, and immune evasion, consistent with a more resistant transcriptional state (Fig. 3B).

**Figure 3.**
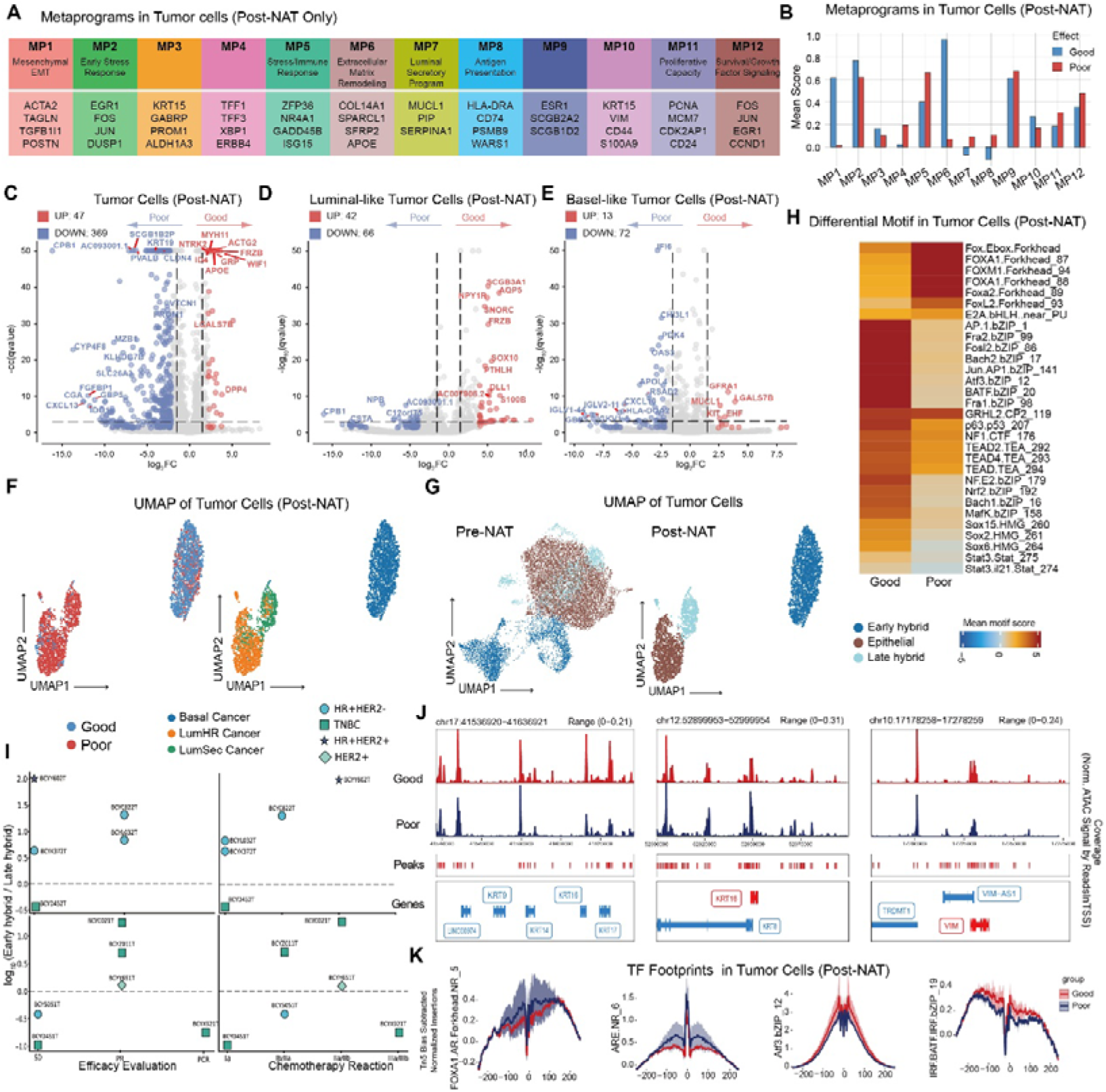
Heterogeneity Analysis of Neoadjuvant Therapy Response in Tumor Cells. (A) Meta programs (MPs) detected from all tumor cells in post-NAT using scRNA-seq data. (B) Barplot shows the enrichment score of MPs in good and poor-response groups from post-NAT samples. (C) to (C) Volcano plots display the differential expressed genes identified in all tumor cells (D), luminal-like cells (F), and basal-like cells (E) from post-NAT samples. The left side corresponds to the upregulated genes of the poor-response group and the right side corresponds to those of good-response group. In the differential comparison between the two groups, genes with an absolute Log2 fold change > 2 and a -log10p-value > 3 were designated as significant. (F) UMAP visualization of residual tumor cells from post-NAT samples, colored by response groups (left) and subtypes (right). (G) UMAP visualization of tumor cells from pre-NAT (left) and post-NAT (right) samples, colored by EMT hybrid states. (H) Heatmap shows differential motifs of tumor cells between good- and poor-response groups. (I) Dotplots display the relationship between NAT-response and early/late hybrid EMT rate. (J) Pseudobulk scATAC-seq signal tracks illustrating differences in chromatin accessibility profiles around the transcription start sites (TSS) of representative EMT marker genes between good- and poor-response post-NAT samples. (K) Transcription factor foot prints in tumor cells from good- and poor-response groups.

We performed differential expression and pathway enrichment analyses on residual tumor cells from both groups(Fig. 3C). In the good-response group, upregulated genes included MYH11 and ACTG2 (smooth muscle contraction/ differentiation), NTRK2 (neurotrophic signaling), and the Wnt/ECM regulators FRZB and WIF1. Additional genes such as ID4, APOE, LGALS7B, GRP, and DPP4 suggested activation of tumor-suppressive, differentiation, and immune-modulatory programs[60–63]. The poor-response group upregulated epithelial and secretory lineage markers (KRT19, SCGB1B2P, CLDN4), immune checkpoint and inflammation-related genes (VTCN1, IDO1, CXCL13, GBP5), stemness/progenitor markers (PROM1, MZB1), and metabolic regulators (CYP4F8, SLC26A3, FGFBP1)[64–67] (Fig. 3C). Collectively, good-response tumors were characterized by activation of EMT, G2-M checkpoint, and tumor-suppressive pathways, whereas poor-response tumors exhibited epithelial, stemness, and immunosuppressive programs (Supplementary Fig. 2F).

Previous research reported that the functional roles of EMT programs vary across different cell types and biological contexts[68]. Thus, we first performed subtype annotation according to canonical epithelial markers. We found that the residual tumor cells in good-response groups were mainly basal cells with highly expression of KRT5, suggesting that EMT signal was largely reflected by an increased proportion of basal cells (Fig. 3F). In contrast, the residual tumor cells in good-response groups contained both luminal and basal cells with highly expression of KRT17 and KRT18 (Fig. 3F). We next performed subtype annotation according to EMT activation. Based on the combined expression of EPCAM, KRT8, KRT14, and VIM, we applied the following classification criteria: Epithelial state: EPCAM+/KRT14+/VIM-; Early hybrid EMT state: EPCAM-/KRT14+/VIM+; Late hybrid EMT state: EPCAM-/KRT14-/KRT8+/VIM+; Late full EMT state: EPCAM-/KRT14-/KRT8-/VIM+[69]. We found that basal cells in the good-response group were in the early hybrid EMT state, consistent with the upregulation of EMT gene sets and increased chromatin accessibility (Fig. 3G and 3J). Late hybrid EMT cells were predominantly derived from the poor-response group (Fig. 3G). We further identified EMT states of epithelial cells in pre-NAT samples and found that in the TNBC subtype, BCYD451T had a lower proportion of early hybrid EMT cells and showed a poorer NAT response (Fig. 3G and 3I). Cells in early hybrid EMT typically retained epithelial adhesion and polarity while acquiring mesenchymal traits, reflecting high plasticity and adaptability. By contrast, late hybrid EMT cells lied closer to a mesenchymal endpoint, characterized by cytoskeletal remodeling, ECM interactions, and activation of migratory pathways, suggesting enhanced invasive/metastatic potential and possible therapeutic resistance[69–70]. Our findings suggested that EMT programs in the good-response group may be associated with wound-healing and tissue-regeneration, whereas those in the poor-response group may be associated with invasive/metastatic potential.

We next excluded confounding effects from subtype composition and analyzed luminal and basal cells separately (Fig. 3D and 3E). In luminal cells, the good-response group upregulated SNORC, NPY1R, FRZB, SCGB3A1, and AQP5, genes linked to hormone receptor positivity, epithelial differentiation, and suppression of EMT/Wnt signaling, suggesting a less invasive and more therapy-sensitive state[71–72]. Conversely, luminal cells in the poor-response group upregulated CPB1, NPB, and AC093001.1, which may reflect aberrant differentiation, neuroendocrine-like features, and heightened resistance potential[73–74] (Fig. 3D). In basal cells, the good-response group exhibited upregulation of genes associated with differentiation, signaling, and epithelial homeostasis (GFRA1, LGALS7B, MUC11, KIT, and EHF), indicating greater differentiation capacity and tissue stability[75–76]. Poor-response basal-like cells upregulated IFI6, CHI3L1, PDK4, OAS3, APOL4, RSAD2, CXCL10, and HLA-DQA2, genes enriched in antiviral responses, inflammation, and metabolic stress, suggesting a heightened immune-activated state[77–78] (Fig. 3E). ScATAC-seq further highlighted differences in motif and transcription factor binding activity between good- and poor-response groups (Fig. 3H and 3K). Poor-response tumors showed enrichment of motifs in the Fox family (FOXA1, FOXM1, FOXA2, FOXL2) and E-box factors (E2A.bHLH), indicative of regulatory programs promoting progenitor-like identity, proliferation, and stemness[45,79–80]. In contrast, good-response tumors exhibited enrichment of AP-1/bZIP family motifs (Fra1, Fra2, Fosl2, Jun, ATF3, BATF, Bach1/2, Nrf2, MafK) as well as p63, NF1, and Sox family (Sox15, Sox2, Sox6) consistent with transcriptional programs supporting epithelial differentiation, tissue homeostasis, antioxidant responses, and immune regulation[39,81–82]. Footprint analysis further revealed that poor-response tumors displayed stronger footprinting of FOXA1 (Forkhead) and ARE (NR) motifs, indicating reliance on Forkhead/nuclear receptor pathways to sustain progenitor-like and hormone-regulated states. Conversely, good-response tumors exhibited stronger footprinting of ATF3 (bZIP) and IRF/BATF (bZIP) motifs, reflecting enhanced AP-1/bZIP and IRF activity associated with stress responses, immune regulation, and differentiation. Footprints reinforced a model in which poor-response tumors are characterized by Forkhead-driven stemness/proliferative networks, while good-response tumors rely on AP-1/IRF-mediated differentiation and immune-modulatory programs. Together, these findings suggest that poor-response tumors are driven by Fox/Forkhead-centered progenitor-like and proliferative programs, whereas good-response tumors are characterized by AP-1-, Sox-, and IRF/BATF-associated networks, together with epithelial lineage factors such as p63 and GRHL2, supporting differentiation, tissue repair, and immune adaptation.

### Characterization of T Cells and Their Clonal Expansion After NAT

To assess differences in the composition of T, NK, and NKT cells across NAT, we integrated T cell transcriptomes from all samples and identified six CD8+ T cell subsets, five CD4+ T cell subsets, one NK cell subset, one NKT cell subset, and one cycling T cell subset (Fig. 4A and 4B). By integrating TCR sequencing with transcriptomic data and performing abundance analysis with miloR, we observed a significant enrichment of early and late terminally exhausted T cells (tTex-early and tTex-late) in post-NAT samples (Fig. 4C). Further inspection revealed that this enrichment was almost entirely driven by the BCYD452T sample (Supplementary Fig. 3A). Upon exclusion of this sample, the abundance of T cell subsets showed no significant differences between pre- and post-NAT (Fig. 4C), suggesting that NAT does not universally induce terminal exhaustion but rather reflects patient-specific effects.

**Figure 4.**
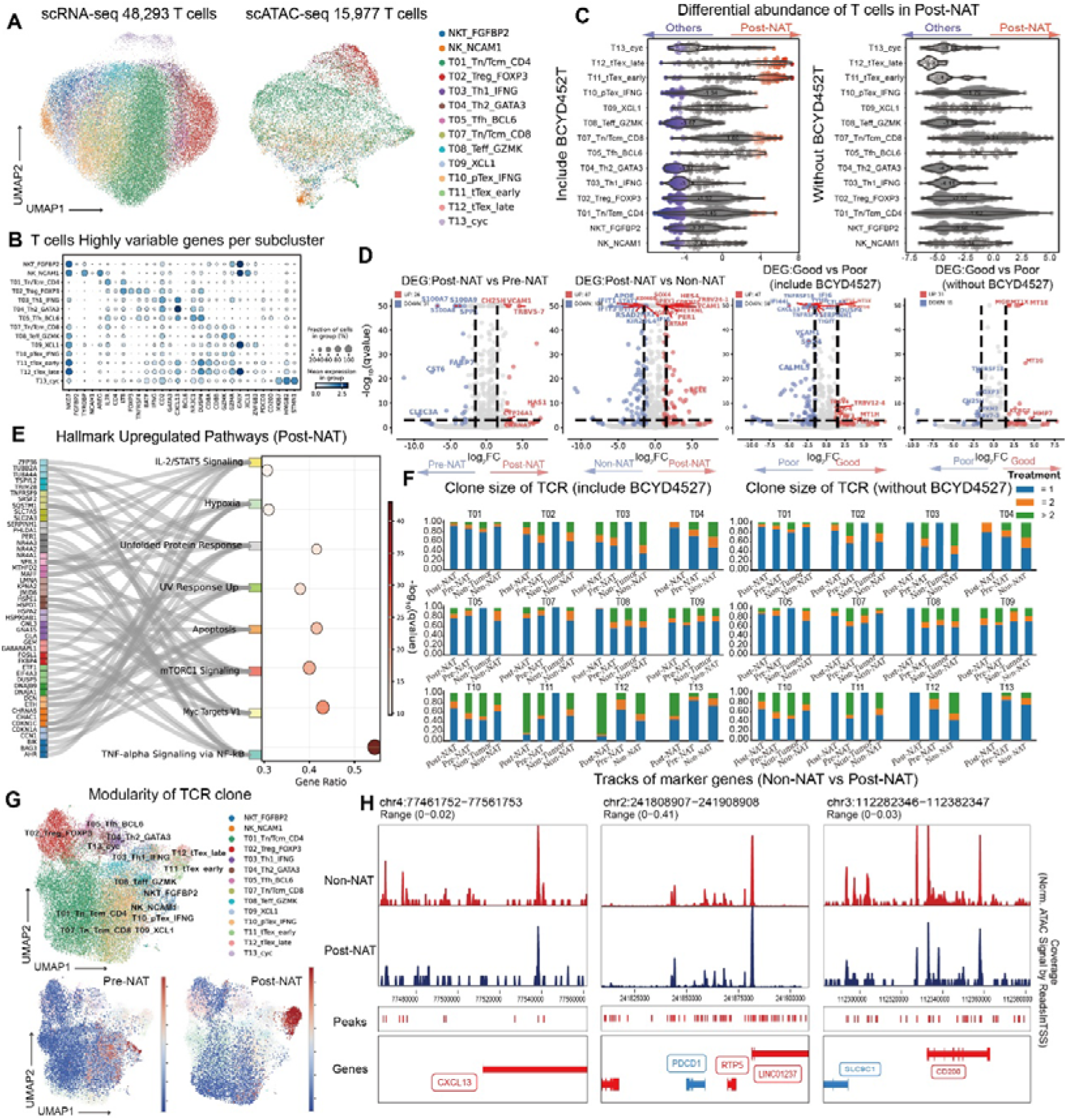
Characterization of T Cells and Their Clonal Expansion After NAT. (A) UMAP visualization of T cells in scRNA-seq (left) and scATAC-seq (right) data. (B) Dotplot heatmap shows highly variable genes in subclusters of T-cells. (C) Beeswarm plots show the log fold change distribution across T cells in scRNA-seq data using post-NAT including BCYD452T vs. other samples (left) and post-NAT without BCYD452T vs. other samples (right). (D) Volcano plots display the differential expressed genes of T cells identified by differential analysis using pre-NAT vs. post-NAT, non-NAT vs. post-NAT, good-response vs. poor-response, and good-response vs. poor-response without BCYD452T. (E) The combination of Sankey and bubble plot depicts significantly upregulated hallmark pathways in T cells following NAT. Each dot represents a hallmark pathway: the x-axis indicates the normalized enrichment score (NES), and the y-axis lists pathway names. Dot color intensity corresponds to the -loglJlJ(FDR) (false discovery rate) value. Dot size reflects the number of genes enriched in the pathway. Gray lines connect pathways to their contributing genes (listed on the left), illustrating gene-pathway associations. (F) Stacked bar plots display the clone size composition of TCRs across T-cell subtypes and treatment conditions in all samples including BCYD452T (left) and excluding BCYD452T (right). (G) UMAP visualization of the modularity of TCR clone in pre- and post-NAT. (H) Pseudobulk scATAC-seq signal tracks illustrating differences in chromatin accessibility profiles around the transcription start sites (TSS) of PDCD1, CXCL13, and CD200 between post-NAT and non-NAT samples.

In scRNA-seq data, T cells in post-NAT samples exhibited marked niche remodeling. TCR variable genes such as TRBV5-7 and TRBV24-1 were elevated in T cells from post-NAT samples, indicating enhanced clonal expansion under antigenic pressure[83]. Moreover, T cells in post-NAT breast tumors showed transcriptional programs consistent with activation-induced differentiation and fate control (KDM6B), together with fine-tuning of cell-cycle progression and activation thresholds (CDKN1C, SPRY1). We also observed expression of molecules associated with tissue localization/retention (CRTAM, GPR183)[84–85] (Fig. 3D).

In post-NAT samples, T cells showed reduced expression of STAT1 together with canonical interferon-stimulated genes (IFIT1, IFIT2, MX1, RSAD2), consistent with attenuated interferon-response transcriptional activity. Decreased CISH is compatible with reduced cytokine/TCR negative feedback. Within the limits of differential expression alone, this pattern may suggest a lower interferon-driven stress tone and a state in which T cells exhibit stronger activation per unit cytokine/TCR input. (Fig. 4D)[86–88].

Pathway enrichment analysis reinforced these observations. Post-NAT T cells exhibited upregulation of IL-2/STAT5, mTORC1, and Myc Targets V1. Compared to T cells in pre-NAT and non-NAT samples, together with adaptive programs including hypoxia response, unfolded protein response (UPR), and apoptosis. Enhanced TNF-α/NF-κB signaling further indicated reprogramming of inflammatory and immune-activation axes (Fig. 4E). These pathway signatures suggest that post-NAT T cells not only acquire greater proliferative and effector potential but also exhibit improved adaptability to metabolic stress, hypoxia, and proteostasis challenges[89–91].

To investigate TCR clonal dynamics, we evaluated both clonal expansion and diversity. Clonal expansion was defined as “more than two cells sharing the same clonotype”. tTex-early and tTex-late subsets in post-NAT showed greater clonal expansion, consistent with therapy-induced antigen-specific responses. However, this effect was lost after excluding BCYD452T (Fig. 4F), confirming that clonal amplification was primarily patient-specific. TCR network modularity analysis revealed that post-NAT networks were concentrated within tTex subsets, reflecting highly focused immune responses (Fig. 4G). Spectratype analysis demonstrated narrowed CDR3 length distributions in post-NAT samples, indicative of reduced diversity and clonal focusing (Supplementary Fig. 3B). Moreover, clonal similarity analyses revealed that pre-NAT samples primarily shared clones among T07, T08, and T10. Post-NAT samples showed clonal overlap between T09 and NKT cells (with overlap in T11 and T12 largely derived from the BCYD452T sample) (Supplementary Fig. 3C). This observation suggested that even under luminal NKT cell apoptosis, NKT cells can still engage in clonal expansion and cooperate with T cells to sustain immune activity. Importantly, NKT cells exhibited greater clonal similarity with multiple T cell subsets in responder patients, underscoring their potential role in coordinating T cell functionality and treatment response.

Polyclonally expanded T cells upregulated CXCL13, while non-expanded T cells preferentially expressed JUN and ZFP36L2 (Supplementary Fig. 3D). CXCL13 has dual implications: it promotes tertiary lymphoid structure (TLS) formation and enhanced antigen presentation, thereby supporting antitumor immunity, but its upregulation is also associated with immune cell clustering in contexts lacking adequate antigen presentation or supportive cytokines, potentially leading to dysfunction[92–93]. Notably, CXCL13 expression was enriched in exhausted T cells, predominantly from the poor-response BCYD452T sample, suggesting its association with unfavourable outcomes (Supplementary Fig. 3D). In contrast, JUN upregulation in non-expanded T cells reflected heightened TCR-mediated activation, whereas ZFP36L2 elevation indicated strengthened post-transcriptional regulation, balancing activation with control of excessive expansion and inflammation (Supplementary Fig. 3D)[94–95]. Together, these data supported a dynamic state of activation-adaptation-homeostatic remodeling in post-NAT T cells: enhanced effector potential counterbalanced by regulatory mechanisms to prevent dysfunction.

To further test the association between CXCL13 and NAT response, we performed response-stratified analyses. With BCYD452T included, CXCL13 was upregulated in T cells from the poor-response group but downregulated in the good-response group; upon removal of this sample, differential expression was no longer observed (Fig. 4D). Responder scoring of key immune regulators, including CXCL13, DUSP4, GZMK, and HLA-DQA1, revealed that these genes were already highly expressed in pre-NAT samples (Supplementary Fig. 3E), supporting their potential as predictors of NAT response.

Finally, scATAC-seq analysis confirmed chromatin accessibility patterns at exhaustion-related loci. CXCL13 and CD200 showed greater accessibility in non-NAT samples, while PDCD1 accessibility was higher in post-NAT (Fig. 4H). Stratification by clinical response revealed higher accessibility of CXCL13 and CD200 in poor responders, whereas PDCD1 was slightly more accessible in good responders (Supplementary Fig. 3F). Collectively, these findings further supported that CXCL13 upregulation associated with poor NAT response, highlighting its potential as a biomarker for NAT response.

### Contribution of Myeloid Compartment to Neoadjuvant Therapy Response

We classified myeloid cells across all samples and identified major subpopulations, including macrophages, monocytes, neutrophils, and dendritic cells (DCs). Macrophages could be further subdivided into five functional clusters: Mac1_SPP1, Mac2_CXCL9, Mac3_FOLR2, Mac4_C3, and Mac5_MMP9 (Fig. 5A and 5B). Functional annotation revealed that SPP1+ macrophages (Mac1_SPP1) were strongly associated with an immunosuppressive tumor microenvironment (TME), promoting EMT and metastatic dissemination. Consistent with previous reports, high SPP1 expression was linked to poor prognosis, reduced survival, and resistance to immunotherapy. In contrast, CXCL9+ macrophages (Mac2_CXCL9) were more likely to participate in antitumor immunity and were significantly enriched in patients with increased tumor-infiltrating lymphocytes and those achieving pathological complete response (pCR). Importantly, the CXCL9/SPP1 expression ratio (CS ratio) has been proposed as a critical predictor of prognosis and immunotherapy response, where high CXCL9 combined with low SPP1 expression generally indicates favorable outcomes[96–98]. Using miloR analysis, we found that both Mac1_SPP1 and Mac2_CXCL9 populations were markedly reduced in post-NAT samples, highlighting the profound remodeling of these key macrophage subsets during neoadjuvant therapy (Fig. 5C).

**Figure 5.**
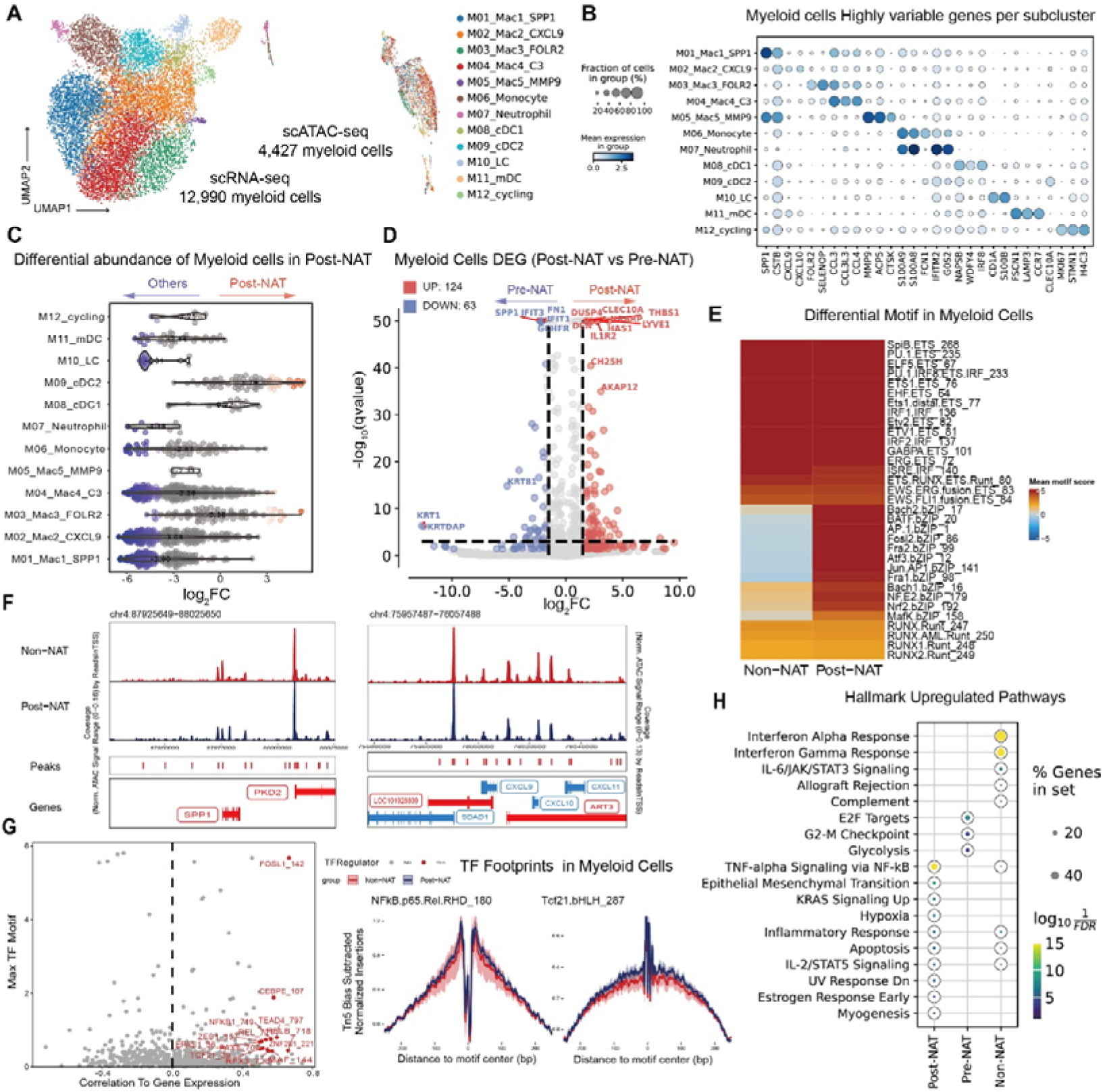
Contribution of Myeloid Compartment to Neoadjuvant Therapy Response. (A) UMAP visualization of myeloid cells in scRNA-seq (left) and scATAC-seq (right) data. (B) Dotplot heatmap shows highly variable genes in subclusters of myeloid cells. (C) Beeswarm plot of the log fold change distribution across myeloid cells using post-NAT vs. other samples in scRNA-seq data. DA neighborhoods at FDR 10% are colored. (D) Volcano plots display the differential expressed genes of myeloid cells identified by differential analysis using pre-NAT vs. post-NAT. (E) Heatmap shows differential motifs of myeloid cells between pre- and post-NAT samples. (F) Pseudobulk scATAC-seq signal tracks illustrating differences in chromatin accessibility profiles around the transcription start sites (TSS) of SPP1 and CXCL9 between post-NAT and non-NAT samples. (G) Scatter plot of transcription factor (TF) motif Enrichment and correlation with gene expression. Each dot represents a TF; dots colored in red highlight TFs with notable enrichment or correlation, with specific TF identifiers labeled. (H) Transcription factor foot prints in myeloid cells from pre- and post-NAT samples. (I) Dot plot displays the enrichment of hallmark biological pathways in Post-NAT, Pre-NAT, and Non-NAT groups. Dot color corresponds to the -log_10_FDR (false discovery rate) value. Dot size represents the percentage of genes in the pathway that are enriched.

Differential gene expression analysis further delineated transcriptional changes in myeloid cells following therapy. Compared with pre-NAT, post-NAT samples upregulated DUSP4, CLEC10A, THBS1, NRARP, LYVE1, DCN, HAS1, IL1R2 and CH25H, while SPP1, IFIT3, FN1, IFIT1 and GCHFR were downregulated (Fig. 5D). These changes indicate that NAT reshaped the functional landscape of myeloid cells: upregulated genes were enriched in immune regulation, ECM remodeling, inflammatory signaling, and lipid metabolism, suggesting enhanced immune activation and tissue-repair potential post-NAT[99–101]; conversely, downregulated genes included mediators of immunosuppression, antiviral responses, and matrix adhesion, implying that therapy may attenuate tumor-promoting and immune-evasive properties of myeloid cells[102–103]. Compared with non-NAT samples, post-NAT samples exhibited increased expression of ADAMTS1, PLEKHG5, ACKR1, and SFRP1, alongside decreased expression of IFIT3, IFIT1, RSAD2, CIRBP, RETN, and CXCL11 (Supplementary Fig. 3G). These transcriptional shifts indicate that post-NAT myeloid cells favor enhanced tissue remodeling and immune-supportive functions, while suppressing excessive interferon signaling and chronic inflammation, thereby contributing to a more favorable TME[104–106].

scATAC-seq motif analysis revealed that post-NAT samples exhibited significantly elevated accessibility of bZIP family motifs compared with non-NAT, including Bach2, BATF, AP-1 (Fosl2, Fra2, Atf3, Jun, Fra1), Bach1, NF-E2, Nrf2, and MafK (Fig. 5E). These transcription factors broadly regulate oxidative stress responses, inflammatory signaling, immune activation, and metabolic reprogramming. Notably, the AP-1 complex plays a central role in macrophage activation and immune signaling, whereas Nrf2, MafK, and Bach1/Bach2 are critical regulators of antioxidant defense and redox metabolism[107–109]. Collectively, these findings suggested that NAT promoted a shift of myeloid cells toward a more stress-adaptive and immunoactive state, potentially enhancing their contribution to therapeutic efficacy.

We further compared the chromatin accessibility of SPP1 and CXCL9 in scATAC-seq data and found that both loci were more accessible in untreated samples (Fig. 5F), consistent with their opposing roles in immunosuppression versus antitumor immunity. ArchR analysis of positive regulatory factors identified FOSL1, CEBPE, NFKB1, TEAD4, REL, RELB, ZEB1, EPAS1, PAX8, ZNF281, TCF21, RFX3, and MAF as significantly upregulated in post-NAT compared to Non-NAT samples (Fig. 5G). These factors encompass multiple key signaling pathways: FOSL1 and NF-κB family members (NFKB1, REL, RELB) regulate inflammatory signaling and immune activation; CEBPE, TEAD4, MAF, and ZEB1 are associated with myeloid differentiation, tissue remodeling, and microenvironmental reprogramming; EPAS1 (HIF2α) and PAX8 reflect hypoxia adaptation and metabolic regulation; and ZNF281, TCF21, and RFX3 suggest remodeling of transcriptional and developmental programs[110–111]. Together, these results indicated that NAT not only directly impacts tumor cells but also profoundly rewires the transcriptional landscape of myeloid cells, enhancing their immune-regulatory and tissue-adaptive potential. Footprint analysis of SPP1- and CXCL9-associated regulators further validated the motif enrichment results.

Pathway enrichment analysis revealed striking differences among the three groups (Fig. 5H). Non-NAT samples were enriched for Interferon Alpha/Gamma Response, IL-6/JAK/STAT3 Signaling, Allograft Rejection, Complement, Inflammatory Response, Apoptosis, and IL-2/STAT5 Signaling, indicating a cytokine-driven, interferon- and complement-dominated inflammatory milieu in untreated tumors. Pre-NAT samples were characterized by enrichment of E2F Targets, G2-M Checkpoint, and Glycolysis, reflecting a state of high proliferative and metabolic demand before NAT. Post-NAT samples showed enrichment of TNF-α Signaling via NF-κB, Epithelial–Mesenchymal Transition, KRAS Signaling Up, Hypoxia, Inflammatory Response, Apoptosis, IL-2/STAT5 Signaling, UV Response Dn, Estrogen Response Early, and Myogenesis, consistent with stress- and adaptation-driven reprogramming after NAT. This pattern highlights not only the reinforcement of immune and inflammatory pathways (TNF-α/NF-κB, IL-2/STAT5, Apoptosis), but also transcriptional shifts toward mesenchymal plasticity and tissue remodeling (EMT, Myogenesis), along with hypoxia adaptation, KRAS signaling, and early estrogen response[112–114]. Collectively, NAT reshaped the TME from a cell cycle– and glycolysis-driven proliferative state (pre-NAT) toward a program characterized by immune–inflammatory activation with EMT and hypoxia adaptation (post-NAT), while non-NAT tumors remained dominated by broad interferon and complement signaling.

Taken together, these findings delineate a trajectory of myeloid cell reprogramming under NAT, whereby neoadjuvant therapy not only targets tumor cells but also fundamentally reshapes the transcriptional and epigenetic states of myeloid cells. This shift, from an immunosuppressive and tumor-supportive phenotype toward one enriched in inflammatory activation, antigen presentation, and immune regulation, likely plays a pivotal role in coordinating immune clearance, fostering adaptive immunity, and stabilizing the post-NAT TME.

### NAT Remodeled Intercellular Communications Within the YBC Tumor Microenvironment

We further resolved the B-cell and stromal compartments. Four major B cell subtypes were identified: naïve B cells (TCL1A+), memory B cells (MS4A1+IGHG1+), T-like B cells (CD3D+), and plasma-like B cells (MZB1+XBP1+) (Supplementary Fig. 4A). Stromal cells were broadly categorized into fibroblasts, endothelial cells, and perivascular-like (PVL) cells. Fibroblasts could be further divided into five functional states: myofibroblastic CAFs (myCAF, ACTA2+TAGLN+), inflammatory CAFs (iCAF, CXCL12+CCL2+), extracellular matrix–remodeling CAFs (ECM-CAF, COL1A1+COL3A1+), antigen-presenting CAFs (apCAF, CD74+HLA-DRA+), and mesenchymal stem–like CAFs (MSC-like CAF, THY1+MGP+) (Supplementary Fig. 4B). Notably, apCAFs were almost exclusively enriched in post-NAT samples along with the increments of iCAFs and ECM-CAFs, suggesting chemotherapy promotes fibroblast reprogramming toward immune-regulatory and ECM-remodeling phenotypes (Supplementary Fig. 4C). Endothelial cells also exhibited marked heterogeneity, comprising six subtypes: venous (Endo Vas-venous, ACKR1+), capillary (Endo Vas-capillary, RGCC+CD36+), arterial (Endo Vas-arterial, IGFBP3+), lymphatic (Endo Lymphatic, LYVE1+), pericyte-like (Endo Pericyte, ACTA2+COL17A1+), and CAF-like endothelial cells (Endo CAF, COL1A1+) (Supplementary Fig. 4D). Among them, multiple endothelial subtypes increased in frequency following NAT, with Endo Vas-venous showing the most prominent rise. PVL cells were classified into four groups, including two pericyte subsets (one characterized by CSPG4+CD44+ expression, and another additionally expressing high CCL2), vascular smooth muscle cells (VSMCs; TAGLN+CNN1+MYL9+), and mesenchymal-like PVL cells (MGP+COL1A1+) (Supplementary Fig. 4E).

To assess how NAT remodeled intercellular communication, we mapped ligand–receptor interactions between tumor epithelial cells and T cells, myeloid cells, B cells, plasma cells, mast cells, and stromal cells across the three sample groups. Basal-like and Luminal-like epithelial cells were analyzed separately, and only ligand–receptor pairs shared by both lineages were retained (Fig. 6A). In epithelial–immune cell interactions, immunosuppressive axes such as SEMA4D–PTPRC and HLA-F–LILRB1 were enriched in non-NAT and pre-NAT samples but diminished after NAT, suggesting attenuation of these inhibitory pathways[115]. CSF1–CSF1R signaling was markedly enhanced post-NAT, indicating a reshaping of myeloid-associated immune axes[116]. Intriguingly, A clear reversal was observed in TNFSF10–TNFRSF10B (TRAIL–DR5) signaling: basal cells had higher activity and luminal cells lower activity in non-NAT and pre-NAT samples, whereas post-NAT samples showed decreased basal cell activity and a strong enhancement of luminal cell activity. This suggested that NAT not only reduces basal-driven immune evasion but also promotes a more pronounced death receptor response in the luminal lineage (Fig. 6A). Meanwhile, angiogenic pathways such as VEGFA/VEGFB–NRP1/NRP2 showed only modest differences overall, but with a trend toward reduced activity in post-NAT samples, consistent with a potential anti-angiogenic effect of chemotherapy.

**Figure 6.**
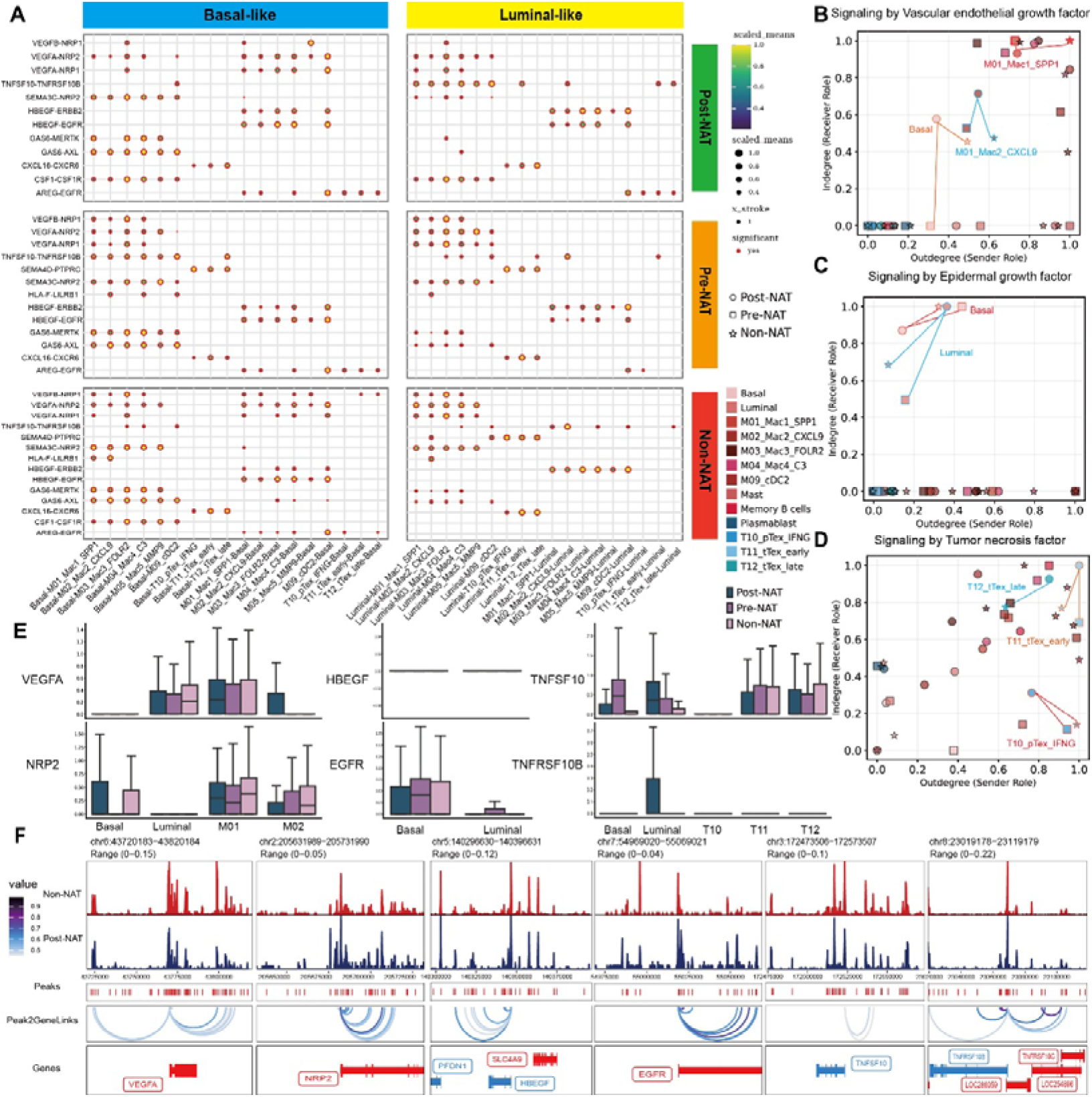
NAT remodeled intercellular communications within the YBC tumor microenvironment. (A) The bubble heatmap shows selected ligand-receptor pairs related to immune checkpoints for interactions of basal-like and luminal-like tumor cells with immune myeloid and T cells in non-, pre-, and post-NAT groups. Dot size indicates the p-value generated by the permutation test, and color indicates the scaled mean expression of each ligand-receptor pair. (B) to (D) Dot plots illustrate the sender (outdegree, x-axis) and receiver (indegree, y-axis) roles of different cell types in signaling via three pathways: vascular endothelial growth factor (B), epidermal growth factor (C), and tumor necrosis factor (D). These plots depict cells from post-NAT (circle), pre-NAT (square), and non-NAT (star) groups. (E) Box plot displays the expression levels of genes involved in vascular endothelial growth factor (VEGFA, NRP2), epidermal growth factor (HBEGF, EGFR), and tumor necrosis factor (TNFSF10, TNFRSF10B) signaling pathways. Subplots show gene expression in cell populations (basal, luminal, M01, M02, T10, T11, T12) across post-NAT, pre-NAT, and non-NAT groups. (F) Pseudobulk scATAC-seq signal tracks illustrating differences in chromatin accessibility profiles around the transcription start sites (TSS) of VEGFA, NRP2, HBEGF, EGFR, TNFSF10, and TNFSF10B between post-NAT and non-NAT samples.

We validated ligand and receptor in scRNA-seq data to support the network analysis. In the VEGF pathway, basal-like tumor cells showed an increase in signaling indegree after NAT, whereas Luminal-like tumor cells exhibited only mild changes in signaling outflow, and their indegree remained near zero. The signaling indegree and outdegree of SPP1+ macrophages (M01_Mac1_SPP1) declined overall, while CXCL9+ macrophages (M02_Mac2_CXCL9) displayed the opposite trend (Fig. 6B). Specifically, the expression of VEGFA arose mainly from macrophages, whereas the receptor NRP2 was enriched in basal cells and a subset of macrophages and was nearly absent in Luminal cells. SPP1+ macrophages (M01) and CXCL9+ macrophages (M02) maintained intermediate-to-high VEGFA expression after NAT, while NRP2 increased in basal cells post-NAT and remained near zero in luminal cells, consistent with enhanced myeloid–basal signaling interactions and limited Luminal receptivity in the network analysis (Fig. 6E). This pattern indicated that NAT suppressed the network activity of immunosuppressive and pro-angiogenic macrophages while enhancing interactions between basal cells and immune-activated or chemotactic macrophages.

In the EGF pathway, basal cells displayed decreased signaling input and output post-NAT, and luminal cells showed increased activity (Fig. 6C). This suggested that the EGFR-dependent survival and repair program shifts from basal to Luminal lineages under chemotherapy pressure. Specifically, HBEGF expression rose post-NAT in luminal cells and in a subset of macrophages, whereas EGFR decreased overall in basal and luminal cells, with a larger drop in basal. Ligand–receptor analysis indicated enhanced Luminal outward and autocrine HBEGF–EGFR interactions post-NAT, while basal outward interactions weakened, corroborating a network-inferred reprogramming in which the EGFR-dependent hub shifted from basal to luminal (Fig. 6E).

In the TNF/TRAIL pathway, both tTex-early and tTex-late T cell subsets showed increased indegree and outdegree, whereas the pTex-IFNG population displayed reduced output but increased input (Fig. 6D). This pattern suggested that exhausted T cells acquired a more central signaling role within the TRAIL axis after NAT. Meanwhile, precursor effector T cells progressively lost signaling capacity. They also become more susceptible to TRAIL–DR5 regulation, which indicates a trajectory toward exhaustion. Specifically, the ligand TNFSF10 increased post-NAT in Luminal cells and exhausted T cells, and the receptor TNFRSF10B was upregulated in tTex-early, tTex-late, and Luminal cells. Accordingly, bidirectional TNFSF10–TNFRSF10B interactions between tTex and epithelial cells strengthened post-NAT. By contrast, pTex_IFNG did not upregulate the ligand but increased receptor expression, indicating a shift in the TRAIL axis from signal sender to principal recipient (Fig. 6E).

In scATAC-seq data, we observed treatment-specific remodeling. In non-NAT samples, VEGFA showed higher accessibility. In post-NAT samples, accessibility increased at NRP2, HBEGF, EGFR, and TNFRSF10B, whereas TNFSF10 was more accessible in non-NAT (Fig. 6F). These patterns agree with the transcriptomic and network results and confirm treatment-induced epigenetic change. Functionally, higher VEGFA accessibility in non-NAT suggests a ligand-ready, pro-angiogenic microenvironment. Increased NRP2 accessibility post-NAT aligns with greater basal receptivity in the VEGF axis and concentrates information flow between myeloid and basal cells. Coordinated opening at HBEGF and EGFR post-NAT provides an epigenetic basis for heightened luminal activity in the EGF pathway, consistent with a shift of the hub from basal to luminal cells. Greater TNFSF10 accessibility in non-NAT matches baseline ligand supply, while post-NAT gains at TNFRSF10B explain higher incoming and outgoing signaling of tTex and the shift of pTex_IFNG from sender to susceptible target.

The network centrality analyses show that NAT rewires tumor-cell signaling dependencies and reconfigures the network centrality of macrophage and T cell subsets across key pathways, thereby inducing functional reprogramming of intercellular communication and driving extensive reorganization of the breast cancer microenvironment. Concordant multi-omic evidence further reveals that residual tumors after NAT acquire adaptive dependencies on angiogenesis, epithelial repair, and immune regulation, highlighting targetable microenvironmental vulnerabilities and potential therapeutic windows.

We compared ligand–receptor interactions across three treatment states, and profiled basal and luminal subsets within the same specimens(Supplementary Fig. 5). Overall, the post-NAT interactome displayed a global enhancement centered on the ECM–integrin axis together with multiple survival/developmental pathways: relative to pre-NAT and non-NAT, FN1–integrin α5β1, COL1A1/COL6A1–integrin α1β1, EFNB1–EPHB2, GAS6–AXL, JAG1–NOTCH1/NOTCH3, and VEGFA–NRP1/NRP2 were upregulated, whereas LAMA3–integrin α3β1 and CD47–SIRPA were downregulated. Functionally, upshifts in ECM–integrin signaling may reflect tissue repair and matrix rebuilding after therapy but can also provide adhesion-survival and migratory support, consistent with stress adaptation and potential drug tolerance[117]. Concomitant elevation of AXL/NOTCH/EPH/VEGF pathways aligns with stemness maintenance, angiogenesis, and immune modulation, suggesting adaptive rewiring of residual clones under chemotherapeutic pressure[118–119]. Conversely, decreased CD47–SIRPA weakens the “don’t-eat-me” signal and is theoretically conducive to phagocytic clearance and innate immune reactivation[120]. Thus, the post-NAT network remodeling exhibits a two-sided phenotype—immune-favorable changes coexisting with reinforcement of survival/regeneration circuits—whose net effect likely depends on their relative balance and on subsequent combination strategies.

In post-NAT samples, basal and luminal subsets showed a division-of-labor in sending/receiving preferences. Basal cells were enriched, as senders, for Laminin–integrin (e.g., LAMA3/LAMC1–integrin) and JAG1–NOTCH axes and were more responsive to these cues as receivers, consistent with signaling programs for re-epithelialization/migration and lineage differentiation/immune interaction, indicating a communication landscape dominated by repair–reconstruction[121–122]. Luminal cells preferentially transmitted EGFR (e.g., AREG–EGFR) and CD47–SIRPA signals, with a tendency toward VEGF–KDR/NRP2, delineating a profile of proliferation–survival, immune evasion, and CSC maintenance[123–124]. Considering that basal cells in this cohort predominantly derive from better responders, whereas luminal cells are overrepresented in poorer responders, these communication biases track with clinical response stratification: Basal cells favor matrix-dependent, differentiation/repair-oriented adaptation, whereas luminal cells favor proliferative, immune-evasive, and angiogenic/CSC-supportive adaptation. Collectively, these axes (integrin, AXL, NOTCH, VEGF, EGFR, CD47) likely participate in shaping response heterogeneity and constitute actionable candidates for rational combinations.

## DISCUSSION

In this study, we integrated scRNA-seq and scATAC-seq data to delineate the NAT-induced remodeling of the TME and cancer cell heterogeneity in young Chinese patients with breast cancer (age ≤ 40). Our cellular analysis provided insights into the molecular mechanisms underlying NAT response. Specifically, NAT reshaped the cellular compositions of TME: epithelial cells, T cells, and myeloid cells decreased, whereas endothelial cells, fibroblasts, and perivascular-like (PVL) cells increased in proportion. The former suggested that neoadjuvant therapy partially cleared tumor burden and associated the immune infiltration; the latter pointed to an adaptive shift under therapeutic pressure toward a more supportive, pro-angiogenic, and pro-fibrotic milieu. We further observed that NAT response correlated with the cellular composition of residual tumor cells: samples with better pathological response (grade Ib–IIIb) exhibited preferential reduction of luminal-like cells with retention of basal-like cells, whereas those with poor response (grade Ia) contained a bulk of luminal-like cells.

NAT also induced alterations in the functional states of tumors: the residual tumor cells in post-NAT was enriched for gene programs related to epithelial–mesenchymal transition (EMT) and matrix remodeling. EMT states are typically associated with increased migratory, invasive capacity, and resistance to apoptosis, providing direct evidence for cell-intrinsic mechanisms of treatment resistance[125]. To strengthen the hypothesis that EMT status relates to NAT response, we classified epithelial cells into Epithelial, Early hybrid EMT, and Late hybrid EMT states in pre- and post-NAT samples. In most cases, a higher ratio of Early/Late hybrid EMT cells were associated with better NAT response (Fig. 3K). For example, BCYD451T (classified as TNBC) has markedly fewer Early hybrid EMT than Late hybrid EMT cells and achieved only Ia by postoperative pathological assessment along with a clinical assessment of SD. An exception was BCYX921T (also classified as TNBC): this case has fewer Early hybrid EMT cells but achieved a postoperative pathological assessment of IIIa and a clinical assessment of pCR. This exception may cause by intrinsic subtypes, inter-individual variability, or differences in treatment regimens, requiring further evaluation for developing personalized regimens.

NAT also impacts specific immune cell subsets, with their alterations linked to treatment response. We initially observed an apparent post-NAT expansion of tTex; however, stratification by sample revealed that this signal was largely confined to the TNBC case BCYD452T. Upon excluding this case, a generalized post-NAT tTex expansion was no longer evident, suggesting that poorly responding TNBC may be prone to tTex accumulation. Notably, CXCL13 was highly expressed within the expanded tTex population. On one hand, CXCL13 can promote tertiary lymphoid structure (TLS) formation and enhance antigen presentation, facilitating specific immune responses; on the other hand, its upregulation is also observed in contexts of localized immune cell clustering with insufficient co-stimulation, implying functional constraint[92–93]. Consistent with this, scATAC-seq showed higher chromatin accessibility at CXCL13 and CD200 loci in non-NAT samples relative to post-NAT samples. We therefore propose that CXCL13+ tTex may serve as a prognostic biomarker associated with suboptimal neoadjuvant response, meriting validation in larger prospective cohorts.

By integrating epithelial–stromal and epithelial–immune communication networks, we found that NAT not only reshapes intercellular communication patterns but also exerts distinct remodeling effects across different tumor cells. Specifically, residual basal-like cells upregulated repair and survival pathways such as ECM–Integrin, NRP2–VEGF, and Notch signaling, whereas residual luminal-like cells upregulated proliferative and immune-evasive programs mediated by HBEGF–EGFR and TRAIL–DR5 axes.

In summary, our single-cell multi-omic analysis charts coordinated remodeling of the TME and tumor cells after neoadjuvant therapy, links EMT phenotypes, lineage-dependent communication axes, and response heterogeneity, and proposes actionable, subtype-directed interventions, thereby providing a clear framework for mechanistic validation and clinical translation. The limited number of cases across different clinical subtypes restricts our analysis of subtype-specific characteristics following neoadjuvant therapy. Future work should supplement samples of the Luminal B and TNBC subtypes to conduct a more comprehensive analysis of their specific characteristics.

## Acknowledgments

This work was supported by the National Key Research and Development Program of China (2020YFA0908700 and 2020YFA0908702) and the National Natural Science Foundation of China (62272246).

## Author Contributions

H.L. curated the clinical cohort and biometrical collection. J.L. conducted the clinical investigation and the experimental design. X.H., W.X., F.L., and Z.R. constructed the bioinformatics pipeline. X.H., F.L., and Z.R. analyzed the scRNA-seq data and immune repertoire. X.H. and W.X. analyzed the scATAC-seq data. B.W., B.S., B.Y., S.L., S.L., and H.L. provided materials and preprocessed the clinical data. X.H., W.X., and J.L. prepared the manuscript with input from all authors. H.L. supervised the material provision, whereas J.L. supervised the bioinformatics analysis. All the authors reviewed, read, and approved the article.

## Ethics approval

This retrospective study was approved by the Ethics Committee of Tianjin medical university cancer institute and hospital (E20210145). Informed consent was obtained from all participants in the cohort for all data acquisition.

## Data availability

The raw sequence data reported in this paper have been deposited in the Genome Sequence Archive in the National Genomics Data Center (NGDC)[126], China National Center for Bioinformation / Beijing Institute of Genomics (CNCB)[127], Chinese Academy of Sciences (GSA-Human: HRA012799), which are publicly accessible at https://ngdc.cncb.ac.cn/gsa-human. All data generated or analyzed during this study are available from the corresponding authors upon reasonable request.

## Code availability

Custom code used in this study is available at GitHub: https://github.com/lyotvincent/BCY-NAT-analysis.

## METHODS

### Patient Material, Ethics, and Consent for Publication

This study was approved by the Ethics Committee of Tianjin Medical University Cancer Institute and Hospital. Written informed consent was obtained from all participants. We enrolled treatment-naive young breast cancer patients under the age of 40, as well as patients who exhibited residual tumors following neoadjuvant therapy. Tumor and adjacent normal breast tissues were collected postoperatively, while pre-NAT tumor tissues were obtained via core needle biopsy in cases where residual tissue was limited after therapy. All specimens were collected between January 2023 and June 2024. These samples were subjected to single-cell RNA sequencing (scRNA-seq) and single-cell assay for transposase-accessible chromatin using sequencing (scATAC-seq). Fresh tissue samples obtained from surgery or biopsy were extensively washed in precooled 1× phosphate-buffered saline (PBS, Invitrogen) to remove blood, adipose tissue, and necrosis, followed by immersion in Tissue Storage Solution (MACS, 130-100-008) for downstream dissociation.

### Tissue Dissociation

Tissue samples were enzymatically digested in a buffer containing 0.35% collagenase IV, 2 mg/mL papain, and 120 U/mL DNase I at 37°C with shaking at 100 rpm for 20 minutes to achieve single-cell dissociation. The reaction was quenched with 1× PBS supplemented with 10% fetal bovine serum (FBS, Gibco), followed by mechanical trituration (5–10 pipette strokes). Cell suspensions were filtered through a 30 μm strainer and centrifuged at 300 ×g for 5 minutes at 4°C. The pellet was resuspended in 100 μL of PBS containing 0.04% bovine serum albumin (BSA), treated with 1 mL of 1× red blood cell lysis buffer (MACS, Cat#130-094-183), and incubated at room temperature or on ice for 2–10 minutes. Cells were centrifuged again (300 ×g, 5 minutes), and dead cells were removed using the MACS Dead Cell Removal Kit (Cat#130-090-101). The resulting cell suspension was washed twice with 0.04% BSA/PBS (300 ×g, 3 minutes, 4°C) and finally resuspended in 50 μL of 0.04% BSA/PBS. Cell viability, confirmed by trypan blue staining, exceeded 85%. Cells were counted using an automated counter and adjusted to a concentration of 700–1200 cells/μL for library construction.

### Single-Cell RNA-Seq Library Preparation

Single-cell RNA libraries were prepared using Chromium Single Cell 3’ v2 and 5’ chemistry kits (10x Genomics), including gel beads, multiplex reagents, and chip systems, according to the manufacturer’s instructions. Approximately 10,000 cells per sample were loaded onto a Chromium chip and encapsulated into oil–aqueous phase GEMs (Gel Beads in Emulsion) together with barcoded gel beads. Cell lysis and reverse transcription of mRNA were carried out within each GEM. Variable regions of TCR (α/β chains) and BCR (heavy/light chains) were selectively amplified via targeted PCR. Libraries were prepared using the Chromium V(D)J Library Construction Kit, which incorporates Illumina adapters and dual-index tags. Sequencing was performed on the Illumina NovaSeq 6000 platform with paired-end, dual-index reads optimized to ensure comprehensive coverage of hypervariable CDR3 regions.

### Single-Cell RNA-seq Data Processing

Raw sequencing data were processed using Cell Ranger (v7.2.0, 10x Genomics) to generate FASTQ files and quantify UMI counts. Barcode/UMI filtering, gene expression matrix construction, and alignment to the human reference genome GRCh38 (v3.0.0) were performed using the multi-pipeline. Low-quality cells (UMI count <1,000, gene count <500, mitochondrial content >20%) and low-frequency genes (expressed in <10 cells) were filtered out. Doublets were identified and removed using Scrublet (v0.2.3)[128] with a score threshold of 0.3. Subsequent normalization, identification of highly variable genes (HVGs), neighborhood graph construction, dimensionality reduction, and clustering were carried out using the Scanpy framework (v1.11.0)[129] in Python (v3.10.14). Batch effects were corrected using harmonypy (v0.0.10)[130] with default parameters.

### Identification of Malignant Epithelial Cells in scRNA-seq data

To delineate malignant epithelial cells, copy number variation (CNV) analysis was performed using infercnvpy (v0.6.1)[20]. Epithelial cells from adjacent normal tissues were designated as the reference group. CNV values were standardized (range: –1 to +1), and genome instability scores were defined as the root mean square of CNV signals per cell. After imputation and denoising, CNV heatmaps were generated to facilitate pattern recognition. Epithelial cells were clustered using Leiden algorithm and visualized via UMAP based on their CNV profiles. The top 5% of epithelial cells with the highest genome instability scores were extracted to construct representative average CNV profiles per tumor.

### Differential Abundance Analysis

Differential abundance (DA) analysis was performed to assess shifts in the relative proportions of cell populations or states across age groups. We applied MiloR[18], a statistical framework that defines overlapping neighborhoods on a k-nearest neighbor (KNN) graph. Based on the top 30 batch-corrected principal components (PCs), a KNN graph was constructed using sc.pp.neighbors with k=50 and Euclidean distance. In MiloR, the neighborhood graph was built with k=20 and Gaussian random vectors of length d=30. Ten percent of the graph’s vertices were sampled using the “reducedDim” strategy with “PCA_harmony” as the optimization target. The number of cells per condition was computed for each neighborhood, and DA testing was performed while controlling the spatial false discovery rate (FDR). Neighborhoods exhibiting significant differential abundance (spatial FDR ≤ 10%) were visualized using swarm plots of log-fold changes between young and non-young samples.

### Intratumoral Transcriptional Heterogeneity and Gene Program Analysis

To characterize transcriptional heterogeneity across treatment regimens and therapeutic responses, we applied GeneNMF[31], an advanced non-negative matrix factorization–based tool designed for gene program discovery across multi-sample scRNA-seq data. Analyses were performed on both all tumor samples and those obtained exclusively after neoadjuvant therapy.

### Single-Cell ATAC-Seq Data Processing

Cells with low-quality profiles were filtered based on nuclear fragment counts, retaining only those exceeding a predefined threshold. To distinguish true chromatin accessibility signals from background noise, we computed transcription start site (TSS) enrichment scores—defined as the ratio of read depth within ±5 kb of the TSS to the flanking background region—and retained cells with scores above threshold. Doublets were removed using ArchR’s (v1.0.3) internal doublet detection algorithm[131]. Dimensionality reduction was performed using latent semantic indexing (LSI)[132], followed by Harmony-based batch correction. Cells were clustered based on a shared nearest neighbor (SNN) graph built from LSI-reduced data.

Gene activity score matrices were computed using ArchR. Specifically, we obtained the pre-imputed GeneScore matrix via getMatrixFromProject() and performed matrix imputation using imputeMatrix() (MAGIC). Imputed values were normalized as log2(imputed GeneScore + 1). Gene activity scores were visualized on the low-dimensional embedding using plotEmbedding() as UMAP overlays, and cluster-level summaries were generated where appropriate.

To call peaks from single-cell ATAC-seq data, we first generated pseudo-bulk replicates per cluster or group using addGroupCoverages(). Reproducible peaks were then identified by invoking MACS2 via ArchR’s addReproduciblePeakSet() (e.g., shift = -40, extsize = 80, --nomodel --nolambda), and the merged peak set was added back to the project with addPeakMatrix(). Cluster-specific differential peaks were obtained using getMarkerFeatures() followed by getMarkers() (e.g., FDR < 0.01 and log2FC ≥ 1, Wilcoxon rank-sum test). For targeted subclustering, we repeated the peak-calling procedure to obtain subgroup-specific peak sets as needed.

Motif enrichment analysis was conducted using HOMER[133]. Motif annotations were added with addMotifAnnotations() (HOMER database), and enrichment across marker peak sets was evaluated with peakAnnoEnrichment(); heatmaps were visualized using plotEnrichHeatmap() to highlight transcription factor (TF) families associated with cluster-defining regulatory programs.

We assessed TF activities using two complementary approaches implemented in ArchR: ChromVAR deviation scores and TF footprinting. For ChromVAR, raw insertion counts over the peak set served as input; GC bias–corrected motif deviation z-scores were computed using addDeviationsMatrix() after motif annotation via addMotifAnnotations(). We also derived a motif “activity” matrix analogous to the GeneScore matrix by retrieving the deviation matrix with getMatrixFromProject() and, when needed, imputing with imputeMatrix() followed by log2-transformation as above. For TF footprinting, motif genomic positions were obtained using getPositions(), per-motif footprints were computed with getFootprints(), and Tn5 sequence bias correction and visualization were performed using plotFootprints(normMethod = “Subtract”).

To infer putative cis-regulatory interactions, we computed peak co-accessibility using ArchR’s addCoAccessibility() with a maximum genomic distance window (e.g., maxDist = 1e6, 1 Mb). Co-accessible links were retained using a correlation cut-off of 0.6 at 1-bp resolution and were subsequently integrated with cluster-specific peak sets and TF motif information to prioritize candidate enhancer–promoter pairs and TF–target relationships within each cell state.

### Annotation of scATAC-Seq Cell Types

To annotate cell types within scATAC-seq data, we leveraged paired scRNA-seq data from the same cohort and performed cross-modality integration and label transfer using the Signac[134] R package. Canonical correlation analysis (CCA) was applied to align gene activity scores from scATAC-seq with transcriptomic profiles from scRNA-seq within a shared latent space. Cell type annotations from scRNA-seq were then transferred to corresponding scATAC-seq cells, and label confidence was determined based on classification probability thresholds.

### Identification of Malignant Epithelial Cells in scATAC-seq data

We estimated DNA copy number alterations (CNAs) from scATAC-seq data following the strategy reported by Satpathy et al. (2019)[21], using the publicly available script 08_Run_scCNV_v2.R (https://github.com/GreenleafLab/10x-scATAC-2019). Briefly, we first partitioned the human genome (hg38) into 10 kb bins with the function makeWindows(genome=BSgenome.Hsapiens.UCSC.hg38, blacklist=blacklist), applying the ENCODE hg38 blacklist regions (ENCFF419RSJ.bed) to exclude problematic loci. CNA profiles were then inferred using the scCNA() function with the parameters neighbors=100, LFC=1.5, FDR=0.1, remove=c(“chrM”,”chrY”). This algorithm estimates copy number status by calculating the mean log2 fold-change of ATAC-seq read coverage for each genomic window relative to its 100 GC-matched neighboring bins. To highlight tumor-specific alterations, we further normalized cancer cell CNV scores by subtracting the average score of tumor microenvironment (TME) cells, yielding a “normalized CNV score” for each cancer cell.

### TCR/BCR Repertoire Analysis

Single-cell V(D)J TCR and BCR enriched libraries were processed using the Cell Ranger vdj pipeline (v7.2.0) with the human reference genome GRCh38 (v3.0.0). Processed TCR/BCR data were analyzed using the scirpy Python package (v1.0.0) for quality control, clonotype assignment, and assessment of clonal expansion[135]. Clonotypes were defined by identical paired full-length CDR3 sequences. Cells sharing the same clonotype and present in multiple cells were considered clonally expanded (clone size >1)[136]. To further characterize TCR repertoire diversity and structure, clonotypes were clustered based on CDR3β amino acid sequence similarity using ir.tl.repertoire_clustering with default parameters.

### Cell–Cell Communication Analysis

To infer cell–cell communication among epithelial, immune, and stromal compartments, we applied CellPhoneDB[137]. We also used Omicverse for analysis, which allowed us to identify the dominant senders, receivers, mediators, and influencers in the intercellular communication network[138]. For each intrinsic subtype and immune cell pairing, we extracted ligand–receptor interactions with P < 0.05 and average expression >1. The frequency of such interactions was quantified and visualized.

### Quantification and Statistical Analysis

Statistical analyses were performed using R (v.4.4.1) and Python (v.3.10.14). Differential expression analysis among cell types was performed by Scanpy (v.1.11.0). Correlations among gene expression levels, functional module scores, and cell type fractions were estimated using Spearman’s rank correlation. For all comparisons, p values less than 0.05 were considered statistically significant. Differential expression genes between treatment conditions, responses, and cell clusters were analyzed using Omicverse (v.1.7.7). Pathway enrichment and gene set enrichment analysis (GSEA) were conducted with the Gseapy (v1.1.9).

