## Supplemental Figures,Table 1 and Table 2 for "Single-Cell RNA and ATAC Sequencing Reveal Cellular Heterogeneity and Chromatin Accessibility Dynamics in Young Chinese Breast Cancer Patients across Pre- and Post-Neoadjuvant Therapy"

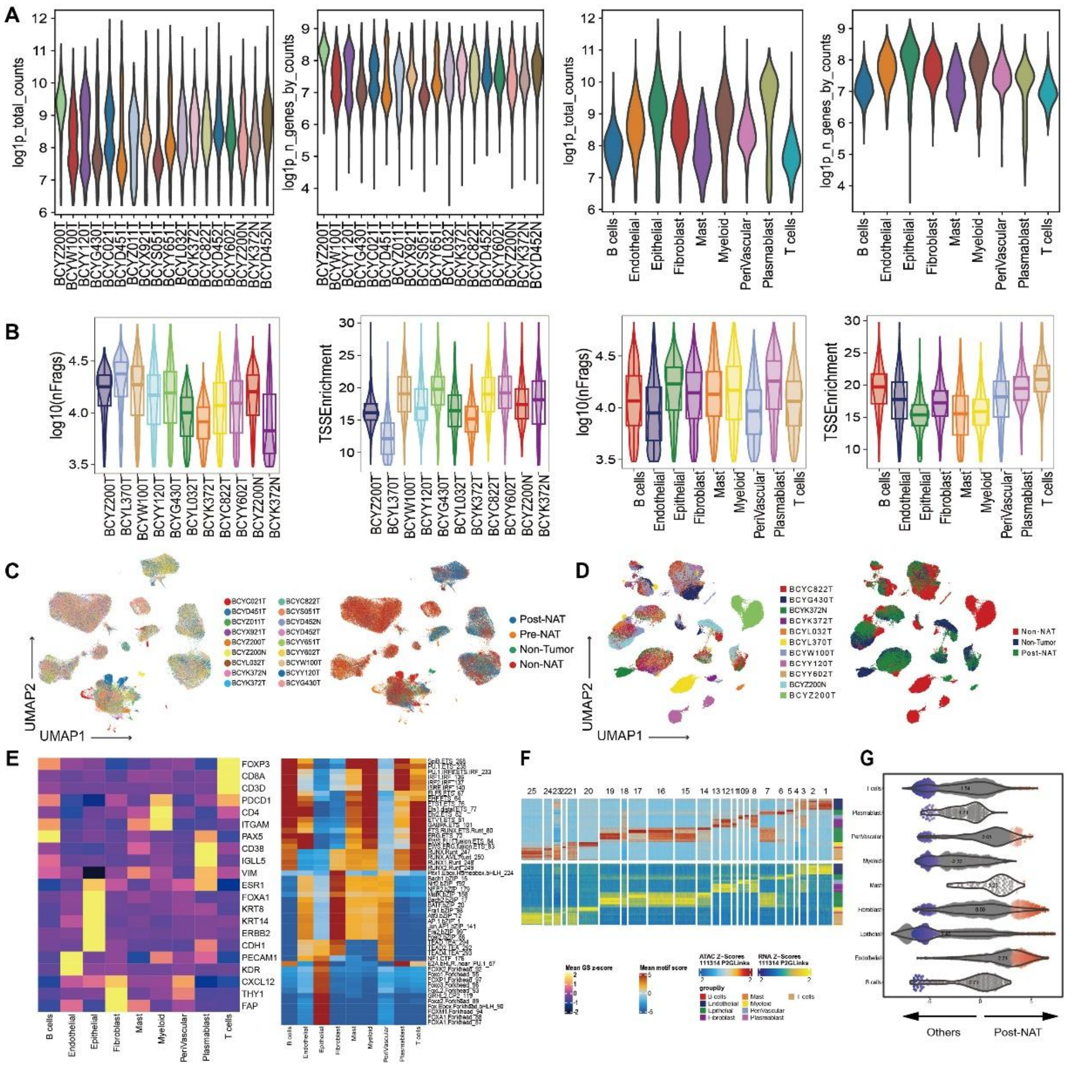


Supplementary Fig. 1 Analysis of quality control and classification of scRNA-seq and scATAC-seq data, related to Figure 1.

(A) Violin plots of quality control metrics for single-cell RNA-seq data. From left to right: log1p_total_counts across samples, log1p_n_genes_by_counts across samples, log1p_total_counts across major cell types, and log1p_n_genes_by_counts across major cell types.

(B) Violin plots showing distributions of scATAC-seq quality control metrics. From left to right: log10-transformed number of fragments per cell (nFrags) across samples, TSS enrichment across samples, log10-transformed number of fragments per cell across major cell types, and TSS enrichment across major cell types.

(C) UMAP visualization of single-cell RNA-seq profiles. The left panel shows cells colored by sample identity, and the right panel shows cells colored by treatment groups.

(D) UMAP visualization of single-cell ATAC-seq profiles. The left panel shows cells colored by sample identity, and the right panel shows cells colored by treatment groups.

(E) The heatmap of gene activity score and motif score of major cell types.

(F) Row-scaled heatmaps of statistically significant distal peak-to-gene links. Each row represents the expression of a gene (bottom) correlated with the accessibility of a distal peak (top).

(G) Beeswarm plot of the log fold change distribution across all cells using post-NAT vs. other samples in scRNA-seq data. DA neighborhoods at FDR 10% are colored.

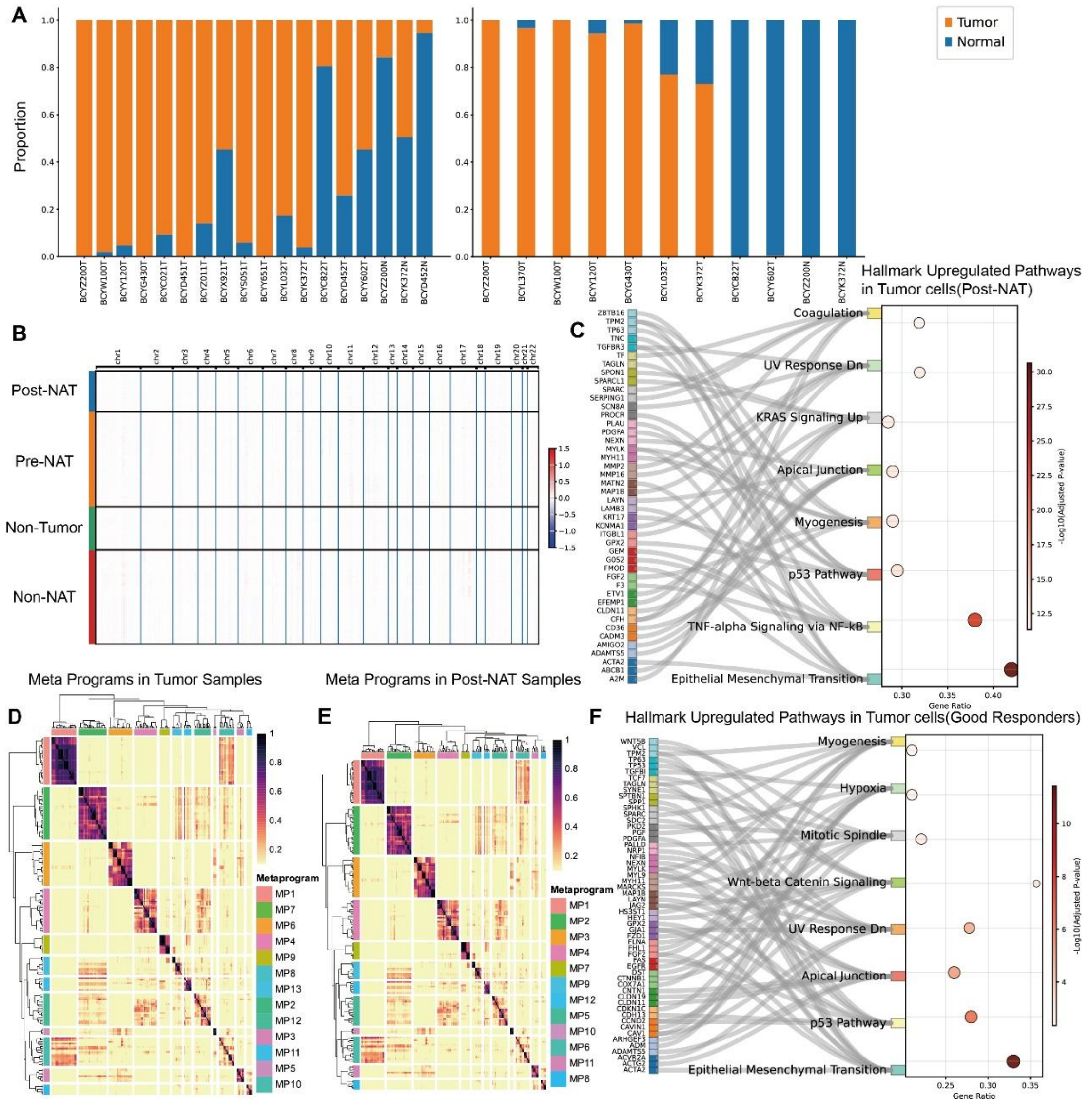


Supplementary Fig. 2 Analysis of copy number variation (CNV), meta-program inference, and pathway enrichment in tumor cells, related to Figures 2 and 3.

(A) Proportions of copy number variation (CNV) states across samples in scRNA-seq and scATAC-seq data.

(B) CNV score heatmap of epithelial cells in scRNA-seq data.

(C) The combination of Sankey and bubble plot depicts significantly upregulated hallmark pathways in tumor cells following NAT. Each dot represents a hallmark pathway: the x-axis indicates the normalized enrichment score (NES), and the y-axis lists pathway names. Dot color intensity corresponds to the $-\log_{10}$(FDR) (false discovery rate) value. Dot size reflects the number of genes enriched in the pathway. Gray lines connect pathways to their contributing genes (listed on the left), illustrating gene-pathway associations.

(D) Heatmap of Meta Programs in all tumor samples.

(E) Heatmap of Meta Programs in post-NAT tumor samples.

(F) The combination of Sankey and bubble plot depicts significantly upregulated hallmark pathways in tumor cells from the good responders. Each dot represents a hallmark pathway: the x-axis indicates the normalized enrichment score (NES), and the y-axis lists pathway names. Dot color intensity corresponds to the $-\log_{10}$(FDR) (false discovery rate) value. Dot size reflects the number of genes enriched in the pathway. Gray lines connect pathways to their contributing genes (listed on the left), illustrating gene-pathway associations.

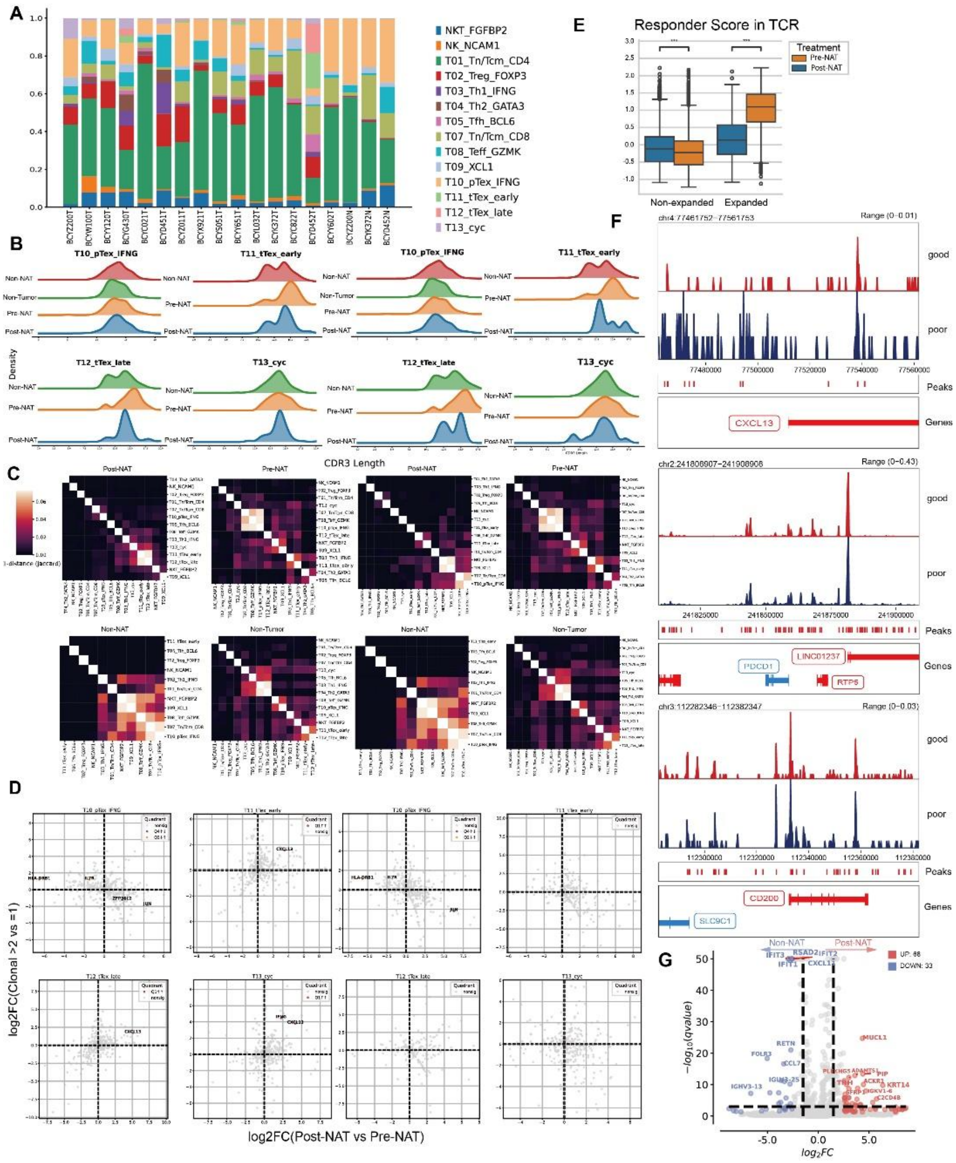


Supplementary Fig. 3 Analysis of T cells and myeloid cells, related to Figures 4 and 5.

(A) Proportions of T cells across samples in scRNA-seq data.

(B) Spectratype analysis of TCRβ CDR3 length distributions. Left panels show results including BCYD452T, while right panels show results excluding BCYD452T.

(C) Clonal similarity heatmaps of TCR repertoires. Left: results including BCYD452T. Right: results excluding BCYD452T. Each four-panel block represents Post-NAT (upper left), Pre-NAT (upper right), Non-NAT (lower left), and Non-Tumor (lower right).

(D) Scatter plots of gene expression changes in expanded versus non-expanded T cell clones. Left: analysis including BCYD452T. Right: analysis excluding BCYD452T.

(E) Box plots of responder scores in TCR repertoires.

(F) Pseudobulk scATAC-seq signal tracks illustrating differences in chromatin accessibility profiles around the transcription start sites (TSS) of PDCD1, CXCL13, and CD200 between good- and poor-response post-NAT samples.

(G) Volcano plots display the differential expressed genes of myeloid cells identified by differential analysis using non-NAT vs. post-NAT.

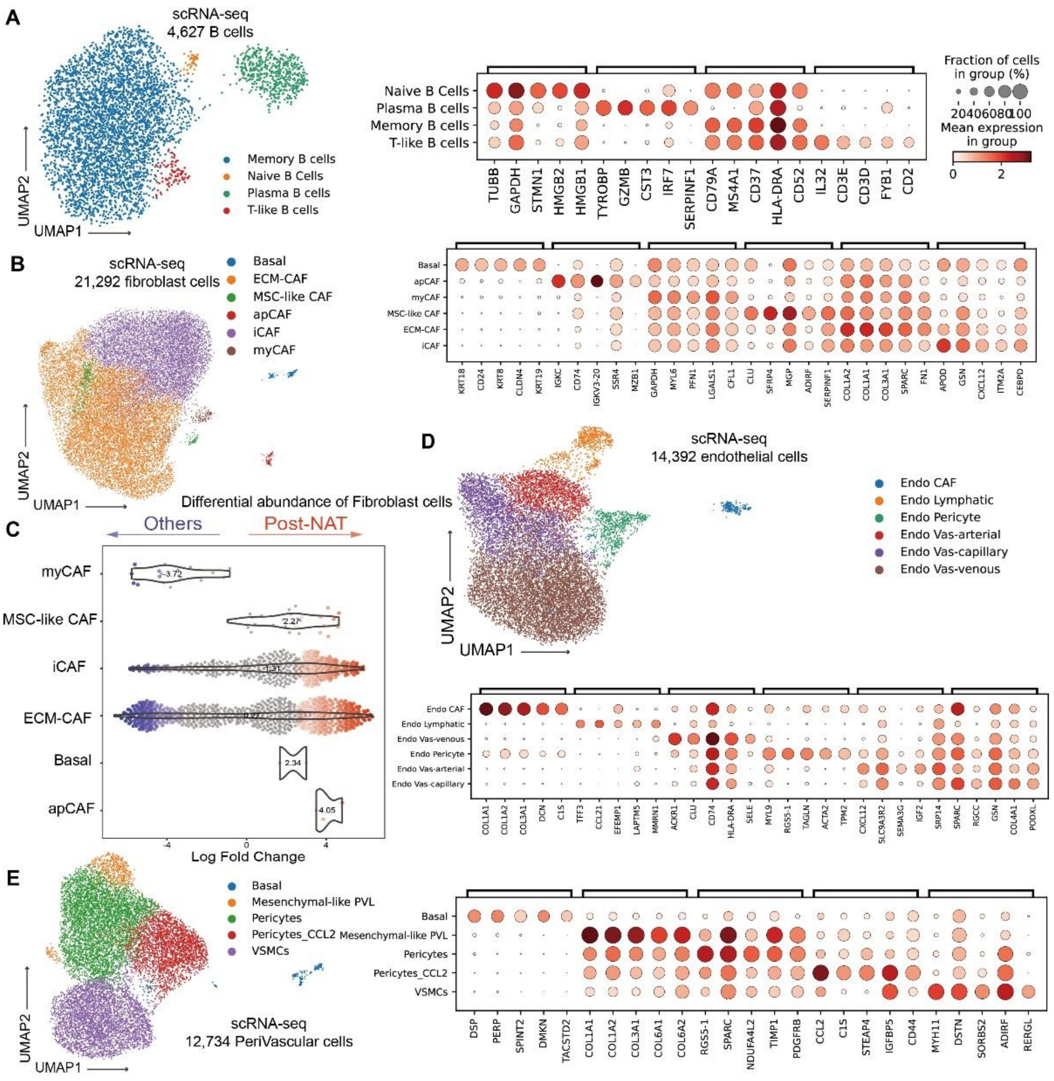


Supplementary Fig. 4 Characterizations of B cells and stromal cells, related to Figure 6.

(A) UMAP visualization of B cells in scRNA-seq (left) and dotplot of marker gene expression in B cell subpopulations (right).

(B) UMAP visualization of fibroblast cells in scRNA-seq (left) and dotplot of marker gene expression in fibroblast cell subpopulations (right).

(C) Beeswarm plot of the log fold change distribution across fibroblast cells using post-NAT vs. other samples in scRNA-seq data. DA neighborhoods at FDR 10% are colored.

(D) UMAP visualization of endothelial cells in scRNA-seq (left) and dotplot of marker gene expression in endothelial cell subpopulations (right).

(E) UMAP visualization of perivascular-like (PVL) cells in scRNA-seq (left) and dotplot of marker gene expression in PVL cell subpopulations (right).

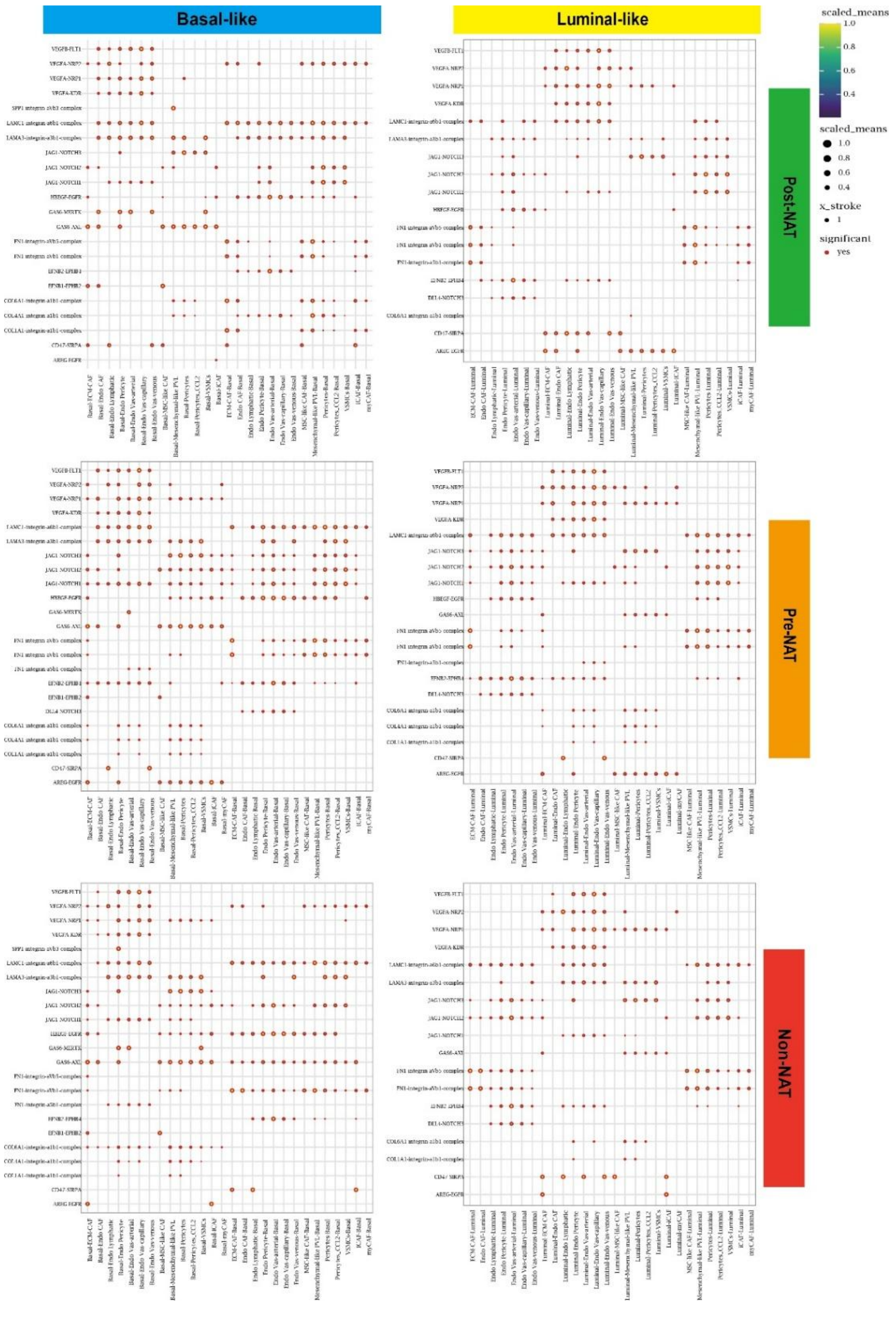


Supplementary Fig. 5 Cell-cell communication analysis of stromal cells and tumor cells, related to Figure 6.

The bubble heatmap shows selected ligand–receptor pairs associated with stromal–epithelial interactions for basal-like and luminal-like tumor cells in non-, pre-, and post-NAT groups. Dot size indicates the p-value generated by the permutation test, and color indicates the scaled mean expression of each ligand–receptor pair.

**Supplementary Table 1. Patient cohort details**

Clinical and pathology details for young breast cancer patients analysed by scRNA-Seq and scATAC-seq in this study.

| PID | Gender | Age | Sampling Methods | scRNA-seq Samples ID | scATAC-seq Samples ID | Gravidity/Parturition | Grade | Cancer Type | ER | PR | HER2 IHC | HER2 ISH (ratio) | Ki67 | P53 | Subtype by IHC | Treatment status | Clinical Stage | Pathology Stage | TNM Stage | BRCA Mutations | Neoadjuvant | Sampling Time | Chemotherapy Reaction | Efficacy Evaluation | Neoadjuvant Treatment Protocol |
|---|---|---|---|---|---|---|---|---|---|---|---|---|---|---|---|---|---|---|---|---|---|---|---|---|---|
| YC02 | Female | 36 | Puncture | BCYC021T | | G0P0 | 3 | IDC | 5 | <1 | 2+ | - | 60 | | TNBC | Naïve | cT2N0M0 | ypT1miN0M0 | II A | | Yes | Before Neoadjuvant | II b | PR | TP + PD1 |
| YD45 | Female | 25 | Puncture | 1T, BCYD452N, BCYD452T | | G0P0 | Unknown | IDC | <1 | <1 | 1+ | | 50 | <1 | TNBC | Naïve | cT2N1M0 | ypT2N1aMx | II B | BRCA1 mutation | Yes | Before Neoadjuvant | I a | SD | EC-T |
| YZ01 | Female | 33 | Puncture | BCYZ011T | | G4P3 | Unknown | IDC | < 1 | < 1 | 0 | | 50 | | TNBC | Naïve | cT2N1M0 | | II B | BRCA1 mutation | Yes | Before Neoadjuvant | I b | PR | TE |
| YX92 | Female | 34 | Puncture | BCYX921T | | G4P2 | 2 | IDC | < 1 | < 1 | 1+ | | 80 | | TNBC | Naïve | cT2N1M0 | ypTisN0Mx | II B | No mutations | Yes | Before Neoadjuvant | III a | PCR | TEC |
| YZ20 | Female | 36 | Surgical | BCYZ200T, BCYZ200N | BCYZ200T, BCYZ200N | G3P2 | 2 | IDC | 90 | 90 | 2+ | - | 50 | 20 | HR+HER2-(Luminal B) | Naïve | cT2N0M0 | pT2N0M0 | II A | | No | | | | |
| YL03 | Female | 36 | Surgical | BCYL032T | BCYL032T | G1P1 | 1~2, 2~3 | IDC | 70 | 60 | 0 | | 30 | 70 | HR+HER2-(Luminal B) | Naïve | cT2N1M0 | ypT2N1aM0 | II B | | Yes | After Neoadjuvant | I a | PR | TE |
| YK37 | Female | 33 | Surgical | BCYK372T, BCYK372N | BCYK372T, BCYK372N | G1P1 | 2 | IDC | 50 | < 1 | 1+ | | 60 | < 1 | HR+HER2-(Luminal B) | Naïve | cT3N0M0 | ypT3N3aM0 | III C | | Yes | After Neoadjuvant | I a | SD | Abiraterone + Leuprolide + Anastrozole |
| YC82 | Female | 33 | Surgical | BCYC822T | BCYC822T | G1P1 | 2 | IDC | 90 | <1 | 0-1+ | | 5 | | HR+HER2-(Luminal B) | Naïve | T2N2M0 | ypT2N0M0 | II A | | Yes | After Neoadjuvant | I b~II a | PR | TE |
| YS05 | Female | 32 | Puncture | BCYS051T | | G2P1 | 2 | IDC | 80 | 30 | 2+ | - | 25 | | HR+HER2-(Luminal B) | Naïve | T2N1M0 | | II B | No mutations | Yes | Before Neoadjuvant | I b | SD | TEC |
| YY65 | Female | 37 | Puncture | BCYY651T | | G1P1 | Unknown | IDC | < 1 | < 1 | 3+ | | 65 | 90 | HR-HER2+ | Naïve | cT2N3M0 | ypT1cN1aMx | III C | | Yes | Before Neoadjuvant | II a | PR | TCbHP |
| YL37 | Female | 39 | Surgical | | BCYL370T | G3P2 | 2~3 | IDC | 70 | 55 | 2+ | - | 30~60 | <10 | HR+HER2-(Luminal B) | Naïve | cT2N1M0 | pT2N1aM0 | II B | | No | | | | |
| YY60 | Female | 37 | Surgical | BCYY602T | BCYY602T | G5P3 | 2 | IDC | 90 | 90 | 3+ | | 15 | 80 | HR+HER2+ | Naïve | cT2N1M0 | ypT3N2aM0 | | | Yes | After Neoadjuvant | II a | SD | TCbHP |
| YW10 | Female | 38 | Surgical | BCYW100T | BCYW100T | G4P1 | 3 | IDC | 85 | 50 | 2+ | + | 35 | 70 | HR+HER2+ | Naïve | cT2N0M0 | T2N0M0 | II A | | No | | | | |
| YY12 | Female | 31 | Surgical | BCYY120T | BCYY120T | G3P1 | 2 | DCIS | 90 | 2 | 3+ | | 25 | 5 | HR+HER2+ | Naïve | cT2N1M0 | pT2(m)N2aM0 | | | No | | | | |
| YG43 | Female | 32 | Surgical | BCYG430T | BCYG430T | G2P1 | 3 | IDC | 2 | 10 | 2+ | + | 70 | <5 | HR+HER2+ | Naïve | cT2N0M0 | T2N0M0 | II A | | No | | | | |

**Supplementary Table 2. Sample details**

Clinical and pathology details for samples analyzed by scRNA-Seq and scATAC-seq in this study.

| Sample ID | Patient ID | Sequencing Technology | Subtype by IHC | Sampling Methods |
|---|---|---|---|---|
| BCYC021T-scRNA | YC02 | scRNA-seq 3' | TNBC | Puncture |
| BCYD451T-scRNA | YD45 | scRNA-seq 3' | TNBC | Puncture |
| BCYZ011T-scRNA | YZ01 | scRNA-seq 5' | TNBC | Puncture |
| BCYX921T-scRNA | YX92 | scRNA-seq 5' | TNBC | Puncture |
| BCYZ200T-scRNA | YZ20 | scRNA-seq 5' | HR+HER2-(Luminal B) | Surgical |
| BCYL032T-scRNA | YL03 | scRNA-seq 5' | HR+HER2-(Luminal B) | Surgical |
| BCYK372T-scRNA | YK37 | scRNA-seq 5' | HR+HER2-(Luminal B) | Surgical |
| BCYC822T-scRNA | YC82 | scRNA-seq 5' | HR+HER2-(Luminal B) | Surgical |
| BCYS051T-scRNA | YS05 | scRNA-seq 5' | HR+HER2-(Luminal B) | Puncture |
| BCYD452T-scRNA | YD45 | scRNA-seq 5' | TNBC | Puncture |
| BCYY651T-scRNA | YY65 | scRNA-seq 5' | HR-HER2+ | Puncture |
| BCYY602T-scRNA | YY60 | scRNA-seq 5' | HR+HER2+ | Surgical |
| BCYW100T-scRNA | YW10 | scRNA-seq 5' | HR+HER2+ | Surgical |
| BCYY120T-scRNA | YY12 | scRNA-seq 5' | HR+HER2+ | Surgical |
| BCYG430T-scRNA | YG43 | scRNA-seq 5' | HR+HER2+ | Surgical |
| BCYZ200N-scRNA | YZ20 | scRNA-seq 5' | Non-tumoral | Surgical |
| BCYK372N-scRNA | YK37 | scRNA-seq 5' | Non-tumoral | Surgical |
| BCYD452N-scRNA | YD45 | scRNA-seq 5' | Non-tumoral | Surgical |
| BCYZ200T-scATAC | YZ20 | scATAC-seq | HR+HER2-(Luminal B) | Surgical |
| BCYL032T-scATAC | YL03 | scATAC-seq | HR+HER2-(Luminal B) | Surgical |
| BCYK372T-scATAC | YK37 | scATAC-seq | HR+HER2-(Luminal B) | Surgical |
| BCYC822T-scATAC | YC82 | scATAC-seq | HR+HER2-(Luminal B) | Surgical |
| BCYL370T-scATAC | YL37 | scATAC-seq | HR+HER2-(Luminal B) | Surgical |
| BCYY602T-scATAC | YY60 | scATAC-seq | HR+HER2+ | Surgical |
| BCYW100T-scATAC | YW10 | scATAC-seq | HR+HER2+ | Surgical |
| BCYY120T-scATAC | YY12 | scATAC-seq | HR+HER2+ | Surgical |
| BCYG430T-scATAC | YG43 | scATAC-seq | HR+HER2+ | Surgical |
| BCYZ200N-scATAC | YZ20 | scATAC-seq | Non-tumoral | Surgical |
| BCYK372N-scATAC | YK37 | scATAC-seq | Non-tumoral | Surgical |
